# A conserved glycan motif overcomes antigenic variation inducing broadly reactive antibodies against the zoonotic pathogen *Streptococcus suis*

**DOI:** 10.1101/2025.06.02.656523

**Authors:** Yao Shi, Göran Widmalm, Charlotte Sorieul, Thomas J. Roodsant, Jeffrey S. Rush, Natalia Korotkova, Antonius A. C. Jacobs, Mirlin Spaninks, Ries Grommen, C. Coral Domínguez-Medina, Irene M. Schimmel, Nicole N. van der Wel, Cameron W. Kenner, Christian Heiss, Parastoo Azadi, Li Tan, Jeroen D. C. Codée, Arjan Stegeman, Constance Schultsz, Lindert Benedictus, Nina M. van Sorge

**Author notes:** Corresponding author: Prof. dr. N.M. van Sorge, Amsterdam UMC location AMC, Department of Medical Microbiology and Infection Prevention, IWO building IA3.211, Meibergdreef 9, 1105 AZ Amsterdam, the Netherlands. These authors have equal contribution to the work (Göran Widmalm, Charlotte Sorieul, Thomas J. Roodsant). These authors jointly supervised the work (Constance Schultsz, Lindert Benedictus, Nina M. van Sorge).

## Abstract

*Streptococcus suis* is a largely neglected but emerging bacterial zoonotic pathogen of global concern for animal welfare, antibiotic resistance development and human health. No effective vaccines are currently available. Here, we identified and characterized the function and structure of two cell wall polysaccharide variants in pathogenic *S. suis* strains using genetic deletion and (heterologous) complementation, lectin staining, glycan composition analysis and specialized NMR spectroscopy. Both glycan variants were anionic polymers that differed in the presence of glucose in the side-chain as a result of allelic variation in a glycosyltransferase gene. Deletion of this variable glycosyltransferase revealed an identical glycan ‘core’ and affected *S. suis* morphology and lysozyme resistance. Immunization of pigs with this core domain induced antibodies recognizing a wide range of antigenically-diverse pathogenic *S. suis* strains. This study provides new insights for developing next-generation glycoconjugate vaccines, whereby a single-glycan target could protect against the emerging zoonotic pathogen *S. suis*.

## Introduction

*Streptococcus suis* is a ubiquitous commensal of the porcine upper respiratory tract that can cause severe infections such as meningitis, septicemia, and arthritis in pigs resulting in high mortality and significant economic losses ^1,2^. In addition to infections in livestock, zoonotic *S. suis* infections are increasingly reported globally and have been identified as the main cause of adult bacterial meningitis several South-East Asian countries ^3–5^. *S. suis* infections are a key driver of antibiotic usage in pigs, thereby contributing to the emergence of multidrug-resistant *S. suis* strains and development of antibiotic resistance in other bacteria ^6,7^. Overall, there is urgent need for innovative preventive measures, particularly vaccines, to prevent *S. suis* infections and improve animal welfare, protect human health, and reduce antibiotic resistance development ^5,8^.

Bacterial surface glycans serve as a protective layer around the bacterium, forming an important physical interface between host and pathogen. Capsular polysaccharides (CPS) form the outermost layer and are important virulence factors, enabling evasion of immune-mediated killing by complement and neutrophils. CPS-based glycoconjugate vaccines induce high functional IgG antibody titers and have been instrumental in reducing mortality and morbidity caused by *Streptococcus pneumoniae* (*S. pneumoniae*), *Haemophilus influenzae* and *Neisseria meningitidis* ^9–11^, especially in young children. However, bacterial pathogens can express a range of structurally-diverse CPS (e.g *S. pneumoniae*), resulting in antigenic variation. Limited cross-serotype protection has significantly offset vaccine-induced reductions in disease incidence as vaccine-targeted serotypes are replaced by other non-vaccine covered serotypes, which can also cause disease^12,13^. In line with human glycoconjugate vaccines, a CPS-conjugate vaccine has been successful in preventing *S. suis* infection in pigs in pre-clinical studies ^14,15^. Similar to human CPS-conjugate vaccines, this vaccine only provided serotype-specific protection, covering only one of the 29 *S. suis* CPS serotypes ^1^. The application of CPS-based vaccines against *S. suis* is further complicated by frequent capsule switching events in the *S. suis* population ^16^, facilitating strain replacement under selective immunological pressure. A new strategy is required to overcome the serotype limitations of traditional CPS-glycoconjugate vaccines, ensuring broad-spectrum protection against *S. suis* infections, support healthy farming, lower antibiotic use in livestock and indirectly reduce zoonotic infections in humans ^16^.

Alongside CPS, Gram-positive bacteria express cell wall-anchored glycan polymers on their surface at high density, which often exhibit less structural variation compared to CPS. Well-studied glycopolymer families include wall teichoic acids (WTA) in staphylococci and rhamnose-rich polysaccharides (RPS) in streptococcal species ^17,18^. Historically, structural variation in RPS composition was employed to classify hemolytic streptococci into Lancefield groups (e.g. Lancefield group A, B, C and G) for diagnosing infections ^17,19^, and to differentiate *Streptococcus mutans* (*S. mutans*) into distinct serotypes (i.e. serotypes c, e, f, k) ^20,21^. However, the role of RPS extends well beyond that of a structural cell wall component and diagnostic agent, fulfilling critical functions in streptococcal cell division and bacterial virulence ^22–26^. Importantly, the conserved nature of the RPS in *Streptococcus pyogenes* (*S. pyogenes*, i.e. group A *Streptococcus*) has led to incorporation of native or modified RPS molecules into multi-component vaccines currently in clinical development ^24,27^.

Similar to *S. pyogenes* and *S. mutans*, we hypothesized that *S. suis* expresses RPS molecules with limited structural variation. Here, we characterized the biosynthesis gene cluster, glycosyl linkage patterns and function of RPS in the zoonotic pathogen *S. suis*. We identified two structural RPS variants in pathogenic *S. suis* lineages, which share a conserved glycan motif and elicit antibodies that recognized a wide range of pathogenic *S. suis* strains in a serotype-independent manner. These findings pave the way for the development of next-generation glycoconjugate vaccines, whereby a single-glycan target could protect against the genetically and antigenically-diverse zoonotic pathogen *S. suis*.

## Results

### Identification of *S. suis* dTDP-rhamnose and RPS biosynthesis gene clusters

The genetic loci encoding the genes required for the biosynthesis of *S. pyogenes* RPS (also known as group A carbohydrate, GAC), and *S. mutans* serotype c RPS (also known as serotype c carbohydrate, SCC) have been identified and characterized previously (Supplementary Fig. 1a) ^22,24^. The putative *S. suis* RPS biosynthesis orthologs *SSU1124* – *1111* form a 14-gene cluster (Fig. 1a), herein designated *srpBCDEGIJKLMNPQR.* This cluster showed a low GC content (Supplementary Fig. 1b), suggesting acquisition by horizontal gene transfer. The genes encode glycosyltransferases (including a rhamnan biosynthesis protein), an ABC transporter, and membrane proteins (Fig. 1a and Supplementary Table 1). In contrast to the GAC and SCC genetic loci ^22,24^, the *S. suis* RPS gene cluster lacked the genes required for dTDP-rhamnose biosynthesis (Supplementary Fig. 1a). However, a 5-gene cluster, which includes putative orthologs of previously identified and characterized dTDP-rhamnose biosynthesis genes *rmlA* – *D* and a gene of unknown function, was identified upstream of *srpBCDEGIJKLMNPQR* in the *S. suis* P1/7 genome (Fig. 1a). We also identified a gene, termed *srpO* (*SSU1672*; Supplementary Table 1), as an ortholog of *S. pyogenes gacO, Streptococcus agalactiae gbcO* and *S. mutans rgpG*, which encode a UDP-*N*-acetylglucosamine (UDP-GlcNAc): Undecaprenol-phosphate GlcNAc-phosphate (GlcNAc-P-P-Und) transferase that catalyzes the first and essential step of GAC, GBC (group B carbohydrate) and SCC biosynthesis, respectively ^23–25^. Similar to other streptococcal species, *srpO* is located separately from the putative RPS gene cluster.

**Fig. 1.**
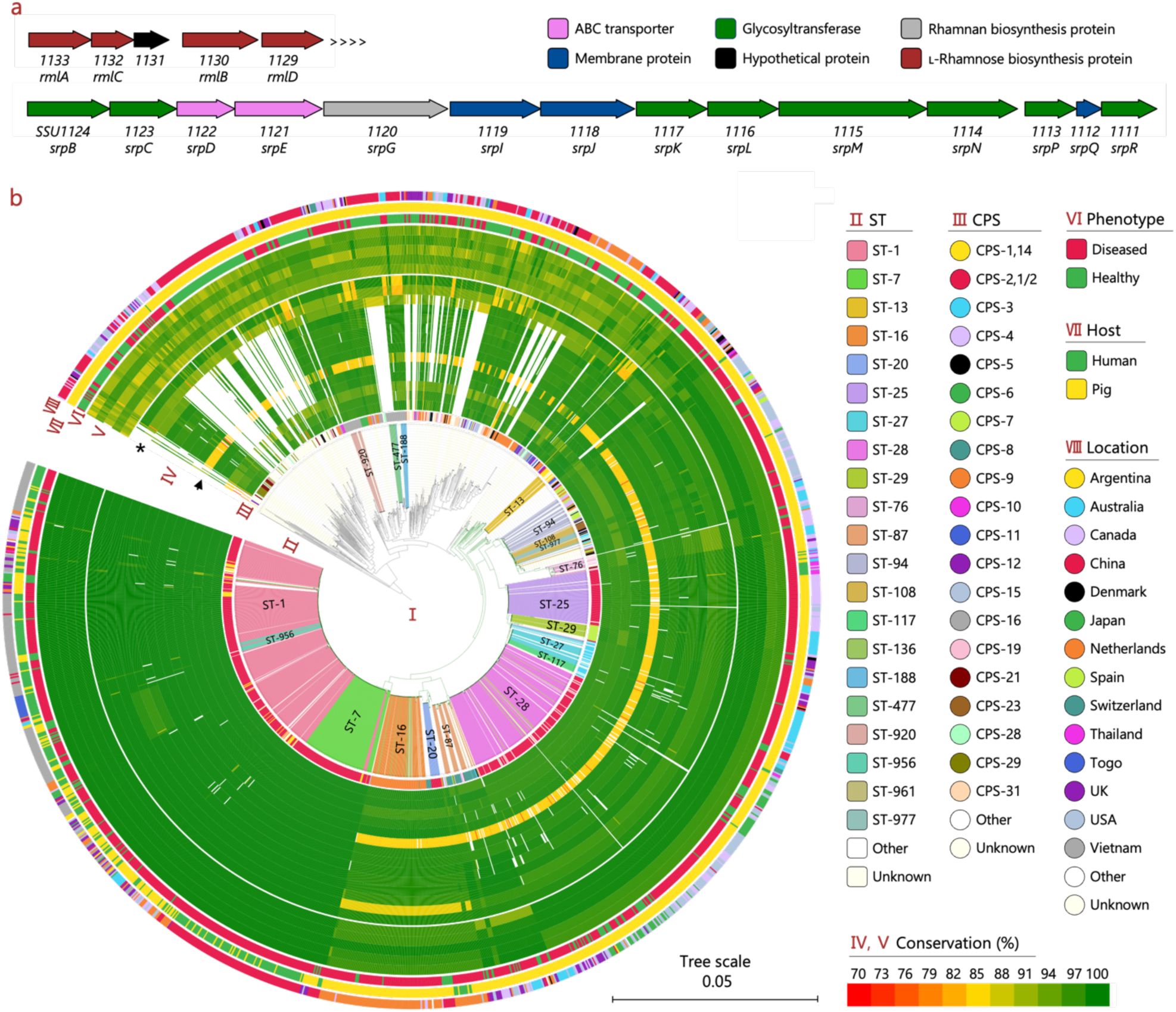
Identification and overview of the RPS biosynthetic gene cluster in the *Streptococcus suis* population. **a**, Schematic representation of the *S. suis* dTDP-rhamnose gene cluster (upper) and RPS gene cluster (lower) in strain P1/7 genome (NCBI accession: NC_012925.1). The arrows are scaled according to gene size, with each predicted function represented by a different color. **b**, Analysis of the putative dTDP-rhamnose and RPS biosynthesis gene cluster in a collection of 1,719 *S. suis* genomes. **I**, Core genome phylogenetic tree. Branches in the pathogenic lineages are colored green. **II**, Sequence type (ST); **III**, serotype-based capsular polysaccharides (CPS) composition. **IV** and **V**, Conservation of RPS (**IV**) and rhamnose (**V**) biosynthesis genes. The dTDP-rhamnose and RPS biosynthesis genes of P1/7 were compared to gene homologues in all other genomes using BLAST. For hits with > 90% coverage and > 80% identity, conservation was calculated by multiplying coverage and identity and visualized using heatmaps. From outer to inner ring of the RPS cluster (**IV**) is *srpBCDEGIJKLMNPQR*, respectively. Asterisk, *srpC*; Arrow, *srpL*. From the outer to inner ring of the dTDP-rhamnose biosynthesis cluster (V) is *rmlA*, *rmlC*, *SSU1131*, *rmlB* and *rmlD*, respectively. **VI**, Disease phenotype of the host. **VII**, Host species; **VIII**, Country where strain was isolated (location).

### Glycosyltransferases SrpC and SrpL show allelic diversity in pathogenic *S. suis* lineages

The level of RPS structural variation differs between species. For example, the GAC biosynthesis gene cluster is highly conserved among the *S. pyogenes* population ^24,28^, whereas the *S. mutans* RPS gene cluster has four identified variants corresponding to its four serotypes ^21,29^. To investigate the conservation of the RPS genes across the genetically-diverse *S. suis* population, we examined the five dTDP-rhamnose biosynthetis genes and the 14 genes of the putative RPS gene cluster by ABRicate in a collection of 1,719 publicly available *S. suis* genomes ^30^. The dTDP-rhamnose biosynthesis genes were ubiquitously present in all genomes and showed little sequence variation (Fig. 1b). In contrast, the composition of the RPS gene cluster varied within the *S. suis* population (Fig. 1b). In the pathogenic *S. suis* lineages, the putative RPS biosynthesis gene cluster was conserved with regard to the number and organization of RPS genes but showed allelic diversity in the two putative glycosyltransferase-encoding genes, *srpC* and *srpL* (Fig. 1b and Supplementary Fig. 2). To gain more insight on gene conservation, we multiplied gene coverage and identity. In the highly pathogenic and zoonotic ST-1 and ST-7 lineages, all RPS genes showed > 91.5% conservation to those in reference strain P1/7. By contrast, the Dutch invasive ST-16 and ST-20 lineages showed < 86.5% conservation in two glycosyltransferase-encoding genes *srpC* and *srpL* compared to P1/7. Sequence variation in *srpL* was also observed in strains belonging to the ST-25 and ST-28 lineages, which are dominant STs in Australia and North America (Fig. 1b). These findings suggest that the RPS glycan composition or structure may differ between pathogenic *S. suis* lineages. Interestingly, RPS genes in non-pathogenic lineages ^16^ had limited homology to those in P1/7 (Fig. 1b and Supplementary Fig. 1c), suggesting structural differences in RPS expressed by pathogenic and non-pathogenic *S. suis* lineages.

To investigate whether *S. suis* strains from different pathogenic lineages express structurally different RPS, we assessed the binding of a panel of plant lectins to ST-1 strain S10 and ST-20 strain 861160. Capsule-deficient mutants were employed to exclude interference from lectin binding to CPS. Soybean agglutinin (SBA) and ricinus communis agglutinin I (RCA I, RCA_120_), which bind to *N*-acetylgalactosamine (GalNAc) and galactose (Gal) with different affinities, demonstrated binding to S10 ΔCPS. In contrast, 861160 ΔCPS was only stained by RCA_120_ (Fig. 2a), suggesting these two strains indeed differ in their RPS composition. To investigate whether structural diversity was linked to the observed allelic diversity in *srpC* and *srpL,* we attempted to delete *srpC* and *srpL* in both strains by homologous recombination. Despite multiple attempts, *srpC* deletion was unsuccessful, suggesting that *srpC* is essential for *S. suis* viability. Deletion of *srpL* in S10 reduced the binding to both SBA and RCA_120_, whereas *srpL* deletion in 861160 increased binding to SBA and RCA_120_ (Fig. 2a). Complementary, plasmid complementation of ΔCPSΔ*srpL* mutants with the homologous *srpL* gene (p*srpL-s*) restored the binding to levels observed in the parent strains, whereas cross-complementation (p*srpL-c*) did not (Fig. 2a). These results suggest that RPS has different compositions in strains S10 and 861160, potentially resulting from genetic differences in *srpL*.

**Fig. 2.**
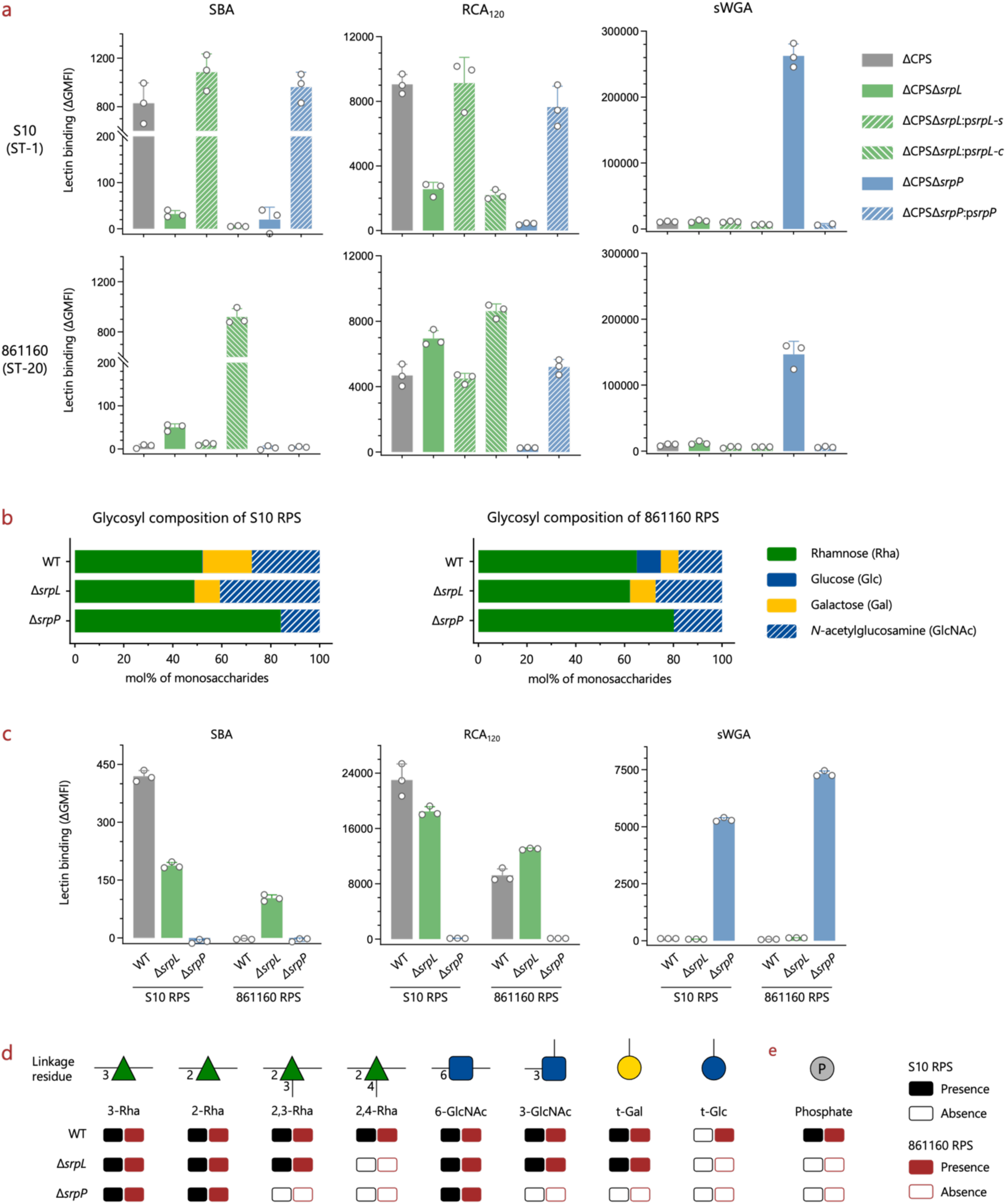
Lectin staining and glycan composition analysis of *S. suis* RPS. **a**, Binding of plant lectins to surface structures of capsular-deficient *S. suis* strains (ΔCPS). Soybean agglutinin (SBA) binds to *N*-acetylgalactosamine (GalNAc) and to a lesser extent galactose (Gal); Ricinus communis agglutinin I (RCA I, RCA_120_) binds to both Gal and GalNAc; Succinylated wheat germ agglutinin (sWGA) has a special affinity to *N*-acetylglucosamine (GlcNAc). Data from biological triplicates of bacteria inoculations are presented as mean values ± SD. ST, sequence type. **b**, Glycosyl composition analysis by GC-MS of TMS derivatives of methyl glycosides of *S. suis* RPS from S10 and 861160 released by mild acid hydrolysis after chemical *N*-acetylation. **c**, Plant lectin binding to isolated RPS from *S. suis* S10 and 861160. Data from three repeats are presented as mean values ± SD. **d** and **e**, Presence and absence of most abundant glycosyl linkage residues (**d**) and phosphate (**e**) of *S. suis* RPS from S10 and 861160. Glycosyl linkage residues were analyzed by GC-MS of partially methylated alditol acetate derivatives. Phosphate was determined by malachite green assay following hydrolysis with hydrochloric acid and digestion with alkaline phosphatase. The original data are shown in Supplementary Table 2 and Supplementary Table 3.

We also deleted *srpP*, a homolog of *S. pyogenes gacI*, which encodes a *N*-acetylglucosamine-phosphate-undecaprenol (GlcNAc-P-Und) synthase, and is essential for biosynthesis of the immunodominant GlcNAc side-chain in *S. pyogenes* GAC ^24,31^. Deletion of *srpP* resulted in loss of both RCA_120_ and SBA binding but conferred binding of succinylated wheat germ agglutinin (sWGA), which has affinity to GlcNAc (Fig. 2a). These observations indicate that *S. suis* RPS contains Gal or GalNAc as terminal moieties and GlcNAc as intermediate moiety. Finally, we attempted to generate a complete RPS-deficient mutant by inactivating *srpO*, but similar to other streptococci, we were unable to obtain mutants after multiple attempts, suggesting that *srpO* is essential to *S. suis*.

### *S. suis* RPS has two structural variants but contains a conserved glycan motif

To gain more detailed insight into the structural composition of *S. suis* RPS, we released RPS from the cell wall of S10 ΔCPS mutant by mild acid hydrolysis after chemical *N*-acetylation, followed by purification by size-exclusion chromatography ^32^. The obtained fraction contained rhamnose, hexose and phosphate (Supplementary Fig. 3a). Further purification of isolated RPS by diethylaminoethyl (DEAE) ion-exchange chromatography showed the presence of DEAE-bound and -unbound fractions (Supplementary Fig. 3b), indicative of a charged and uncharged fraction, respectively. Gas chromatography-mass spectrometry (GC-MS) based glycosyl composition analysis revealed that the two fractions of the S10 wild-type (WT) RPS contained the same monosaccharides, i.e. Rha, Gal and GlcNAc at a molar ratio of 5.2:2.0:2.7 (Supplementary Fig. 3c). By contrast, the 861160 WT RPS, prepared in a similar manner, contained Rha, Glc, Gal and GlcNAc in a molar ratio of 6.5:1.0:0.7:1.8 (Fig. 2b). These results confirm that the S10 WT RPS and the 861160 WT RPS differ in their glycan composition by the absence or presence of Glc, respectively.

Since deletion of *srpL* and *srpP* changed plant lectin binding (Fig. 2a), we performed GC-MS based glycosyl composition analysis on the RPS of these mutant strains. RPS isolated from ΔCPSΔ*srpL* mutants showed decreased Gal content in S10 and complete absence of Glc but no difference in Gal in 861160 (Fig. 2b). The isolated RPS molecules were also probed by plant lectins (Fig. 2c) and the results were consistent with lectin binding to whole bacteria and glycosyl composition analysis. RPS isolated from the S10 Δ*srpP* and 861160 Δ*srpP* deletion strains revealed an identical glycan composition, consisting of Rha and GlcNAc and lacking Gal and Glc (Fig. 2b). Correspondingly, the samples showed identical plant lectin staining profiles (Fig. 2c). Overall, these observations confirm that the RPS glycan composition is different between strain S10 and 861160 and can be attributed to genetic variation in *srpL*.

To investigate the structural differences in RPS from WT and related deletion strains in S10 and 861160, a detailed glycosyl linkage analysis of RPS from S10 and 861160, using GC-MS of partially methylated alditol acetate derivatives, was performed. Both the S10 and 861160 WT RPS contained 3-Rha, 2-Rha, 2,3-Rha, 2,4-Rha, 6-GlcNAc, 3-GlcNAc and terminal Gal as the most abundant linkage residues, whereas the 861160 WT RPS contained additional terminal Glc (Fig. 2d). Deletion of *srpL* resulted in the loss of 2,4-Rha in both strains, and loss of terminal Glc in 861160, in line with results from composition analysis (Fig 2d). Upon deletion of *srpP*, RPS showed the presence of 3-Rha, 2-Rha, and 6-GlcNAc and loss of 2,3-Rha, 2,4-Rha, 3-GlcNAc and terminal Gal in both strains, and the absence of terminal Glc in 861160 (Fig. 2d). Furthermore, NMR spectral data from ^1^H,^13^C-HSQC experiments of mutant strains of *S*. *suis* indicate that Δ*srpP* RPS from both S10 and 861160 contain similar linear polysaccharides consisting of trisaccharide repeating units, whereas RPS of Δ*srpL* mutants additionally contained similar branched side-chains resulting in hexasaccharide repeating units. Thus, RPS of Δ*srpL* mutants represents a conserved glycan motif, which may serve as a good candidate for a broad-spectrum vaccine against *S*. *suis*.

### Glycerol phosphate is linked to different sugars in ST-1 strain S10 and ST-20 strain 861160

GAC and SCC carry a negative charge because of the glycerol phosphate (GroP) modification, which increases susceptibility to cationic antimicrobial peptides (AMPs) and cationic antibacterial proteins like human group IIA-secreted phospholipase A_2_ (hGIIA) ^26,28,32^. Similarly, both *S. suis* WT RPS variants were negatively charged and contained phosphate (Fig. 2e and Supplementary Fig. 3), suggesting *S. suis* RPS also undergoes modification with GroP. However, phosphate was not detected in Δ*srpL* or the Δ*srpP* RPS from both S10 or 861160 (Fig. 2e), implying that the terminal Gal transferred by SrpL S10 or terminal Glc transferred by SrpL 861160 serve as the GroP acceptor. To confirm the presence and location of GroP, purified RPS from both WT strains were analyzed by NMR spectroscopy. The S10 RPS contained a phosphodiester group, as indicated by the NMR chemical shift ^33^, *δ*_P_ 0.71, and a ^1^H,^31^P-HMBC experiment revealed correlations to protons in the spectral region 4.05 – 3.80 ppm (Fig. 3a). For 861160 RPS, the corresponding NMR data were *δ*_P_ 0.86 and 4.20 – 3.80 ppm (Fig. 3b). Furthermore, in the ^13^C NMR spectra, resonances with closely similar chemical shifts were observed, viz., *δ*_C_ 67.20 (*J*_CP_ 5.6 Hz), *δ*_C_ 71.55 (*J*_CP_ 7.6 Hz) and *δ*_C_ 63.02 for the S10 RPS, and *δ*_C_ 67.17 (*J*_CP_ 5.6 Hz), *δ*_C_ 71.56 (*J*_CP_ 7.6 Hz) and *δ*_C_ 63.05 for the 861160 RPS. These NMR findings resemble those from *S. pyogenes* GAC, in which the substituent *sn*-Gro-1-*P* is linked to position 6 of a β-ᴅ-Glc*p*NAc residue as a side-chain in the polysaccharide ^26^. Similarly, in *S. mutans* SCC, *sn*-Gro-1-*P* is also linked to position 6, though in this case to the α-ᴅ-Glc*p* ^29^.

**Fig. 3.**
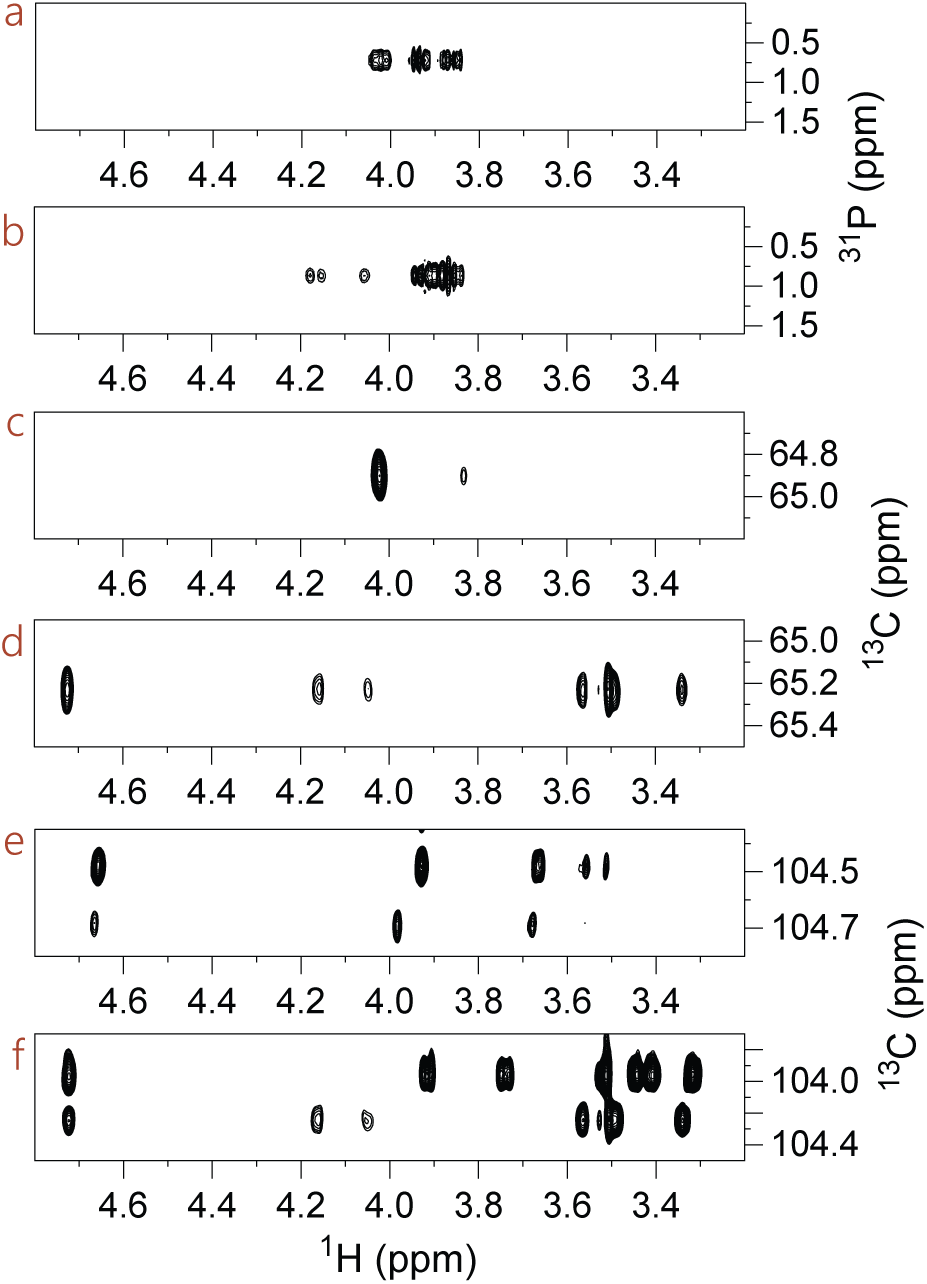
Selected NMR spectral regions of RPS from S10 and 861160. **a** and **b**, ^1^H,^31^P-HMBC NMR spectrum of (**a**) RPS from S10 (ST-1 strain) and (**b**) RPS from 861160 (ST-20 strain), identifying phosphodiester-linked entities. Spectral region from a ^1^H,^13^C-HSQC-TOCSY NMR experiment (mixing time of 200 ms) showing correlations from C6 of a hexopyranose residue substituted by *sn*-Gro-1-*P* at the hydroxymethyl group for (**c**) RPS from S10 revealing only a limited number of cross-peaks consistent with a *galacto*-configuration of the sugar and (**d**) RPS from 861160 displaying cross-peaks for a full ^1^H,^1^H spin-system for a hexose, including the anomeric proton, consistent with a *gluco*-configuration of the sugar. **e**, Spectral region from the ^1^H,^13^C-HSQC-TOCSY NMR experiment for the RPS from S10 showing correlations from anomeric carbons of hexopyranose residues devoid of a GroP substituent (major, *δ*_H1_/*δ*_C1_ 4.66/104.47, ^1^*J*_C1,H1_ 162 Hz) and carrying a GroP substituent (minor, *δ*_H1_/*δ*_C1_ 4.67/104.67, ^1^*J*_C1,H1_ 162 Hz) revealing only a limited number of cross-peaks consistent with a *galacto*-configuration of the sugar and (**f**) the corresponding spectral region from the ^1^H,^13^C-HSQC-TOCSY NMR experiment for the RPS from 861160 displaying cross-peaks for full ^1^H,^1^H spin-systems, including hydroxymethyl protons, of hexopyranose residues devoid of a GroP substituent (major, *δ*_H1_/*δ*_C1_ 4.723/103.96, ^1^*J*_C1,H1_ 165 Hz) and carrying a GroP substituent (minor, *δ*_H1_/*δ*_C1_ 4.725/104.24, ^1^*J*_C1,H1_ 164 Hz) consistent with a *gluco*-configuration of the sugar.

Additional NMR analysis confirmed that both RPS variants contained terminal Gal residues whereas only the 861160 RPS contained terminal glucose residue(s) (vide infra). The ^1^H,^13^C-HSQC NMR spectra of the two RPS structures showed cross-peaks of hydroxymethyl groups shifted to higher ^13^C NMR chemical shifts, indicative of phosphorylation ^34^, with an increase of ca +3 ppm to ∼65 ppm. The ^1^H,^13^C-HSQC-TOCSY NMR spectrum of the S10 RPS showed a cross-peak at *δ*_H5_ 3.83 (Fig. 3c) besides the cross-peak at *δ*_H6_/*δ*_C6_ 4.02/64.90, consistent with a Gal residue having *δ*_C5_ 74.41. The corresponding ^1^H,^13^C-HSQC-TOCSY NMR spectrum of 861160 RPS showed cross-peaks at *δ*_H5_ 3.57 and *δ*_H1_ 4.725 (Fig. 3d) besides cross-peaks at *δ*_H6a_/*δ*_C6_ 4.05/65.23 and *δ*_H6b_/*δ*_C6_ 4.16/65.23, consistent with a Glc residue having *δ*_C5_ 75.58 (*J*_C5,P_ 8.3 Hz). The assignments of Gal and Glc residues in S10 and 861160 RPS, respectively, were consistent with the lower number of cross-peaks in S10 (Fig. 3e) compared to the full ^1^H,^1^H spin-system of the hexose in 861160 RPS (Fig. 3f). Further analysis confirmed that S10 RPS contained a Gal residue substituted at O6 by an *sn*-Gro-1-*P* group. This was supported by the presence of a cross-peak, *δ*_H5_/*δ*_C4_ 3.83/69.02, in a ^1^H,^13^C-H2BC NMR spectrum, indicating a large ^2^*J*_H5,C4_ coupling constant, as observed in Gal and lactose ^35^. The absolute configuration of the substituent is assumed to be *sn*-Gro-1-*P* similar to that found in *S. pyogenes* GAC ^26^. Thus, complementary NMR analysis showed that the RPS of S10 and 861160 carry the GroP substituent at O6 of Gal and Glc residues, respectively, both of which are β-linked, as inferred from the magnitude of their ^1^*J*_C1,H1_ coupling constant (NMR data given in the legend to Fig. 3). The identified *S. suis* RPS biosynthesis gene cluster lacked an apparent ortholog of GroP transferases-encoding *gacH* and *sccH*, which are essential for a transfer of the GroP moieties on GAC and SCC, respectively ^22,26^ (Fig. 1a and Supplementary Fig. 1a). Therefore, the GroP transferase responsible for the GroP modification of *S. suis* RPS is likely located separately from the RPS biosynthesis gene cluster and remains to be identified.

### RPS side-chain is important for *S. suis* biology

To explore the function of the RPS side-chain in *S. suis*, we compared the growth and morphology of S10 Δ*srpL* and Δ*srpP* mutants to S10 WT. The S10 Δ*srpL* mutant, which lacks the Gal-GroP side-chain, showed slower growth in rich culture media (Fig. 4a) and increased chain length (Fig. 4b). By contrast, the Δ*srpP* mutants, that only expressed the RPS trisaccharide backbone, displayed normal growth but significantly increased chain length (Fig. 4b and 4c) and enhanced self-aggregation after overnight culture (Supplementary Fig. 4). Scanning electron microscopy showed that encapsulated Δ*srpP* mutant is cone-shaped, with some swollen, small blebs and fractional cells (Fig. 4c middle column and Supplementary Fig. 5a, 5b), but transmission electron microscopy showed that septa were being formed (Fig. 4c right column). In contrast, all ΔCPS mutants showed smoother surface and the ΔCPSΔ*srpL* and ΔCPSΔ*srpP* mutants showed longer cell length comparing to the parent strain ΔCPS (Supplementary Fig. 5c right column). Complementation of deleted genes restored the growth, chain length and cell length of the mutants, but not the cell shape of the encapsulated Δ*srpP* mutant (Fig. 4a – c and Supplementary Fig. 5c). Altogether, these observations indicate that the RPS side-chain is important for *S. suis* morphology, chaining, and growth.

**Fig. 4.**
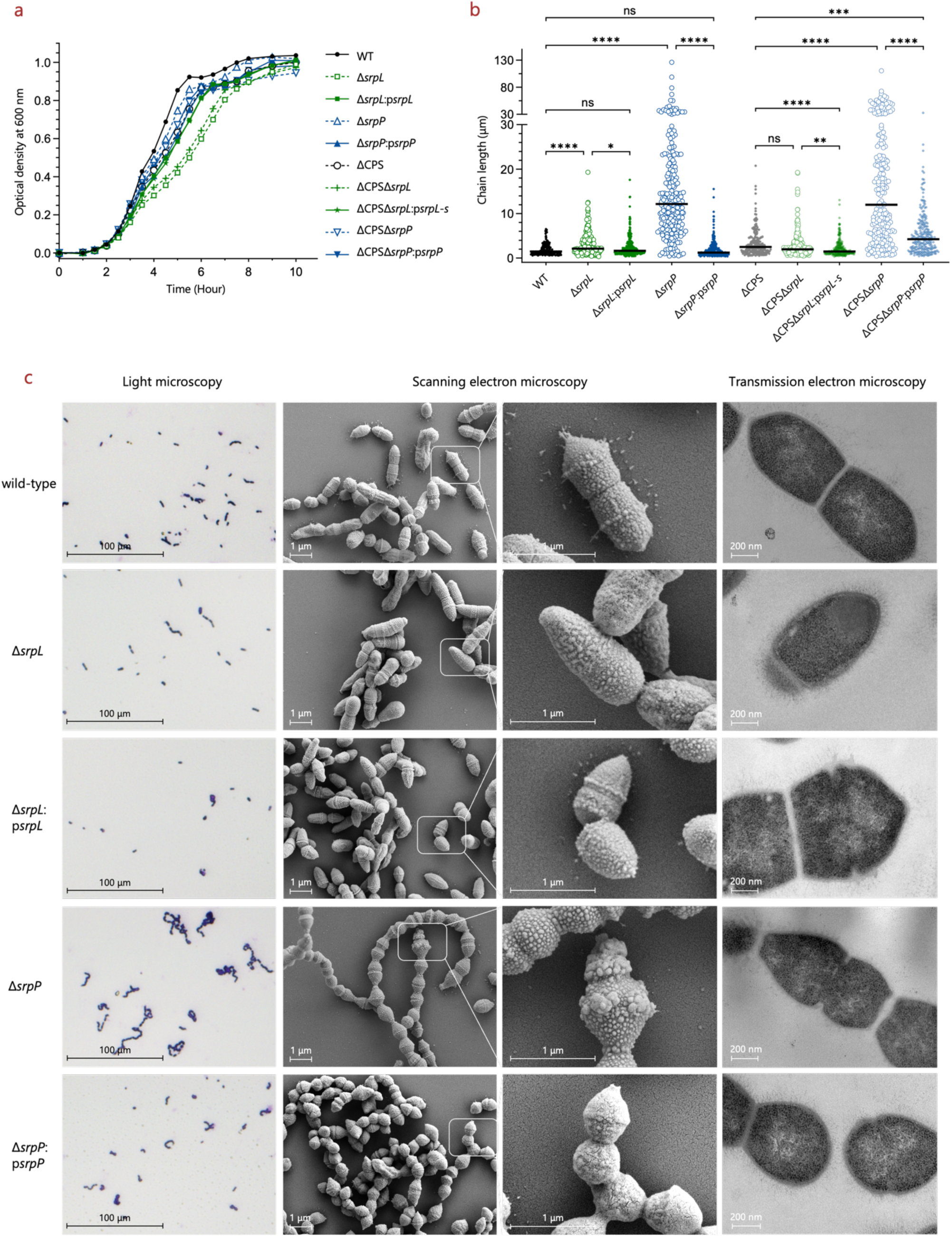
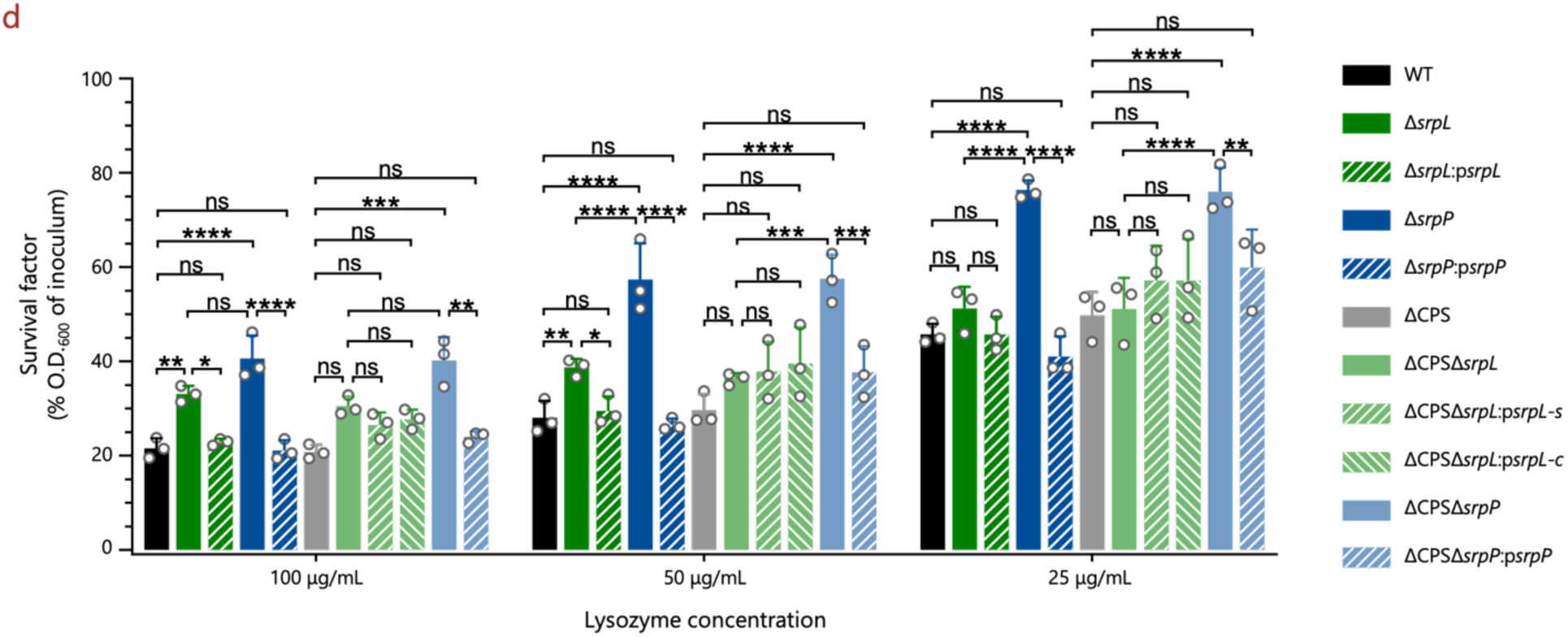
RPS side-chain is important for *S. suis* cell division, morphology and lysozyme resistance. **a**, Growth curves of S10 wild-type and isogenic *srpP* and *srpL* deletion mutants and complementation strains in a capsule expressing or capsule deficient (ΔCPS) background. Bacteria were incubated at 37°C with gentle shaking in test tubes and optical density (O.D.) at 600 nm was measured every 30 min. Data of Δ*srpL* and Δ*srpP* mutants were collected from three and two independent experiments, respectively. Representative curves are shown. **b**, Chain length of S10 wild-type and isogenic *srpP* and *srpL* deletion mutants and complementation strains. Overnight culture in THY with gentle shake was employed for Gram staining (**c** left column) and multiple pictures of every single strain were taken under 40 × microscope objective. The quantification of chain length was performed by measuring all bacterial chains in every single picture using ImageJ, until at least 200 chains were measured. The line represents the median value. *P* values were calculated by Kruskal-Wallis test with Dunn’s multiple comparisons test. \**p*≤0.05; \*\**p*≤0.01; \*\*\**p*≤0.001; \*\*\*\**p*≤0.0001; ns, not significant. **c**, Representative microscopy images of encapsulated S10 wild-type and isogenic *srpP* and *srpL* deletion mutants and complementation strains. Light microscopy images were taken on Gram stained stationary phase bacteria from overnight culture in THY; scanning (middle 2 columns) and transmission (right column) electron microscopy images were taken on exponential phase bacteria (O.D. ≈ 0.4 in THY). **d**, RPS side-chain contributes to *S. suis* lysozyme susceptibility. ΔCPS represents capsule deficient background. Data from biological triplicates of bacteria inoculations are presented as mean values ± SD. *P* values were calculated by two-way ANOVA with Tukey’s multiple comparisons test. \**p*≤0.05; \*\**p*≤0.01; \*\*\**p*≤0.001; \*\*\*\**p*≤0.0001; ns, not significant.

In *S. pyogenes*, the GAC-GlcNAc side-chain and the GroP modification affect the susceptibility of *S. pyogenes* to human antimicrobial defenses, such as antimicrobial peptides, human type IIA-secreted phospholipase A2 (hGIIA) and lysozyme ^24,32,36^. To determine whether specific RPS epitopes impact *S. suis* immune resistance, we assessed lysozyme susceptibility. Both the Δ*srpL* and Δ*srpP* mutants showed increased resistance to lysozyme compared to the isogenic WT strain (Fig. 4d), with mutants lacking the entire RPS side-chain (Δ*srpP* and ΔCPSΔ*srpP*) being most resistant (Fig. 4d). These results indicate that the RPS side-chain contributes to lysozyme susceptibility in *S. suis*, in line with observations in other streptococci.

### Conserved RPS motif is a broad-spectrum vaccine candidate against *S. suis*

Structural analysis of RPS from two pathogenic *S. suis* strains revealed limited structural variation and the presence of a shared glycan antigen consisting of Rha, GlcNAc and Gal. We therefore hypothesized that this shared glycan antigen could elicit broadly reactive antibodies unaffected by subtle glycan variations. To test this, RPS was purified from the 861160 ΔCPSΔ*srpL* mutant and conjugated to the carrier protein CRM_197_ ^37^ using click chemistry (copper(I)-catalyzed azide-alkyne cycloaddition, CuAAC) ^38^ to generate a glycoconjugate construct (Supplementary Fig. 6a). The glycoconjugate was adjuvanted with X-Solve ^39^ and three-week-old piglets were immunized intramuscularly twice at two-week intervals. No increased antibody reactivity to either the carrier protein or RPS was observed on day 14 after the first immunization. However, on day 28, two weeks after the second immunization, seven (58.3%) out of twelve piglets showed IgG responses against the immunizing glycan (Fig. 5a), while all piglets developed IgG against CRM_197_ (Supplementary Fig. 6b). In contrast, no IgG reactivity against the carrier protein or the RPS glycan was detected in the control group (Fig. 5a and Supplementary Fig. 6b).

**Fig. 5.**
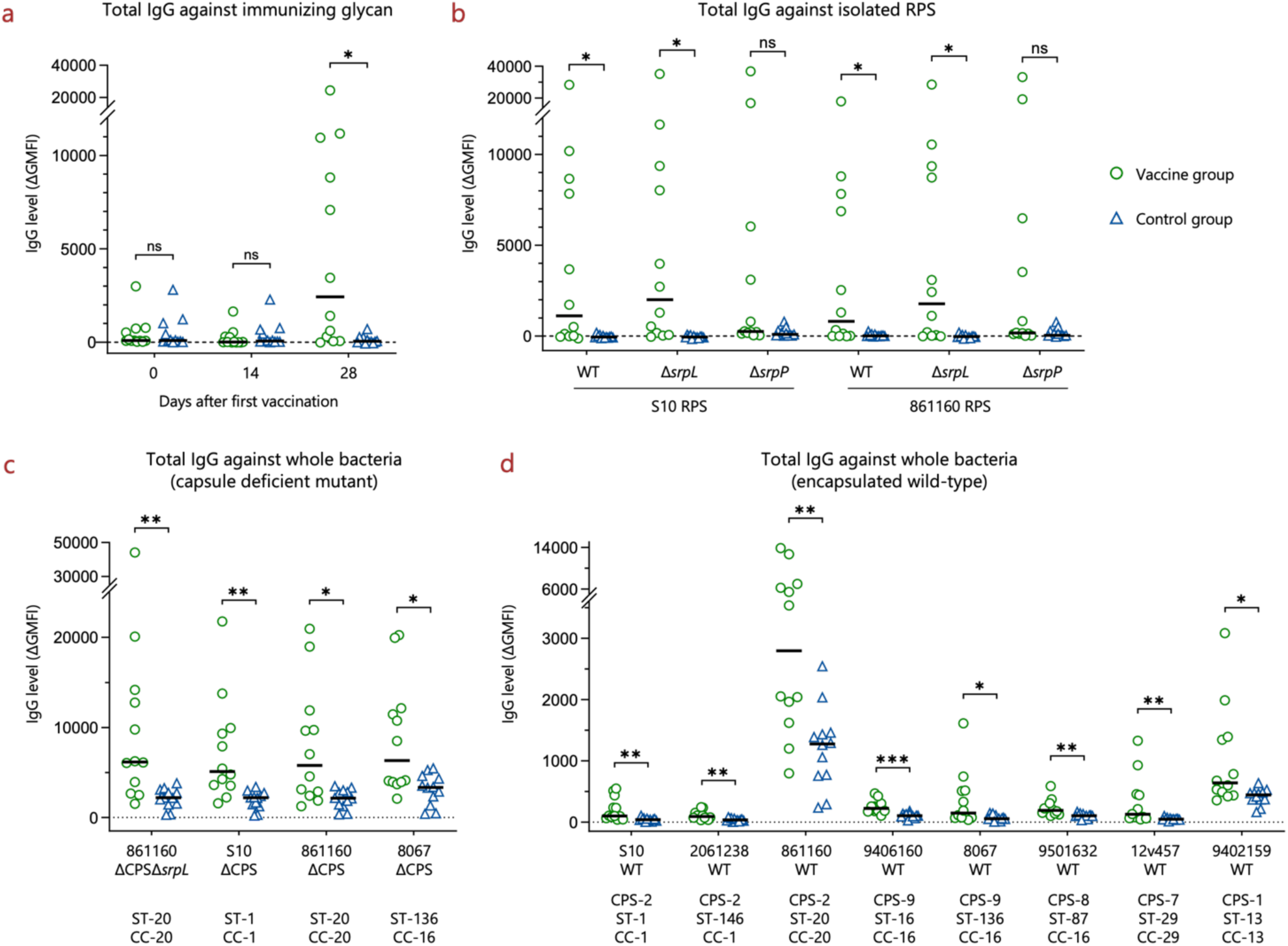
Conserved RPS motif induces IgG antibodies with broad reactivity against *S. suis*. **a**, The conserved RPS structure as isolated from 861160 ΔCPSΔ*srpL* induces IgG in piglets when administrated as a glycoconjugate vaccine on day 28 after two vaccinations. Three-week old piglets (n = 12) were immunized twice using adjuvanted RPS-glycoconjugate, with two weeks interval. Sera were collected on days 0, 14 and 28 and analyzed for IgG responses to the immunizing glycan using RPS-coated magnetic beads. **b**, The conserved RPS structure induced cross-reactive antibodies against different RPS variants including the wild-type structures. Sera from day 28 were used to evaluate cross reaction. RPS from 861160 ΔCPSΔ*srpL* represents the immunizing glycan. The correlations of IgG binding to the immunizing glycan and different RPS variants are shown on Supplementary Fig. 7a. **c** and **d**, The IgG antibodies that are elicited through vaccination with the conserved RPS structure are reactive against both (**c**) CPS-deficient *S. suis* strains and (**d**) encapsulated WT strains from different serotypes and sequence types in the pathogenic lineages. Sera from day 28 were incubated with exponential phase bacteria to evaluate antibody binding. The correlations of IgG binding to the immunizing glycan and different *S. suis* strains are shown on Supplementary Fig. 7b and 7c. All sera were used at a 1:100 dilution; Antibody binding was accessed by flow cytometry. The lines represent the median values. *P* values were calculated by Mann-Whitney test with Holm-Šídák’s multiple comparisons test. \*\*\**p*≤0.001; \*\**p*≤0.01; \**p*≤0.05; ns, not significant. WT, wild-type; CPS, capsule polysaccharide type (serotype); ST, sequence type; CC, clonal complex.

IgG antibodies elicited by the RPS-conjugate were reactive against purified WT, Δ*srpL* and Δ*srp*P RPS of both S10 and 861160 (Fig. 5b). To further investigate whether these antibodies could cross-react with diverse *S. suis* strains, pig sera were incubated with representative *S. suis* strains from various genetic backgrounds within the pathogenic lineages. The RPS-reactive IgG induced by immunization bound to four different CPS-deficient *S. suis* strains (Fig. 5c), as well as eight encapsulated WT strains representing five disease-associated serotypes and eight STs (Fig. 5d). These results indicate that the shared RPS motif can elicit antibodies that recognize a broad range of pathogenic *S. suis* strains and are not hindered by natural RPS variation. This conserved RPS motif warrants further investigation as a potential target for the development of a broad-spectrum vaccine against the diverse zoonotic pathogen *S. suis*.

## Discussion

A broadly protective *S. suis* vaccine would not only prevent disease in pigs but also reduce antibiotic use and lower the risk of human infections through zoonotic transmission. While RPS known as GAC is highly conserved in *S. pyogenes* and explored in clinical vaccine development ^24,27^, the vaccine potential of the conserved glycan motif of the four *S. mutans* RPS variants has not been studied. Here, we provide an initial molecular dissection of *S. suis* RPS, defining the RPS biosynthesis gene cluster and elucidating the glycosyl linkage patterns of the two structural RPS variants in pathogenic *S. suis* lineages. Importantly, immunization of piglets with a glycan motif common between the RPS variants elicited antibodies that were broadly reactive against this genetically diverse pathogen, opening novel avenues for RPS-based *S. suis* vaccines.

We demonstrated that SrpP is essential for side-chain biosynthesis, which is in line with the function of the GacI homolog in *S. pyogenes* ^24,31^. In *S. pyogenes*, GacJ forms a complex with GacI and significantly stimulates its catalytic activity ^31^. As SrpQ contains a similar functional domain as GacJ, we infer that SrpP is a GlcNAc-P-Und producer, which initiates the side-chain biosynthesis supported by SrpQ. There is also an overlap between the GAC and SCC side-chains and the *S. suis* RPS side-chain with regard to function. First of all, we demonstrated that the side-chain of RPS is crucial for *S. suis* growth and morphology as well as resistance to antimicrobial peptides, consistent with findings in *S. pyogenes* and *S. mutans* ^22,24,29,40^. Moreover, the GAC side-chain is considered to be a virulence determinant in multiple *S. pyogenes emm* types ^24,41^. Similarly in *S. suis*, *srpP* was identified to be essential for bacterial survival in the blood, brain and cerebrospinal fluid (CSF) of pigs in a transposon insertion mutagenesis assay ^42^ and upregulated when *S. suis* is cultured in pig blood compared to Todd Hewitt Broth (THB) ^43^. In addition, deletion of *srpP* significantly decreased biofilm formation ^44^. Altogether, the RPS side-chain is likely to be more broadly involved in *S. suis* virulence and pathogenesis.

We successfully conjugated the conserved RPS motif to a carrier protein using click chemistry and used this construct to immunize piglets. Although all piglets showed an IgG response to the carrier protein on day 28, only seven (58%) of 12 piglets developed IgG responses against the immunizing glycan. However, the immune system of piglets is not fully developed until about four-week of age ^45–47^, which may partially explain this incomplete response. In addition, we only attempted a single immunizing strategy and adjuvant, which may not have been optimal for inducing glycan-specific antibodies in piglets ^8^. For implementation of a *S. suis* vaccine in practice, the vaccine would actually be applied to sow, which would then transfer the IgG antibodies to the piglets ^39^. Therefore, future studies should explore alternative immunization conditions and adjuvants to improve glycan-reactive IgG responses.

The IgG antibodies in piglets that did respond to vaccination, did not only bind to the conserved glycan motif but also recognized the two glycan variants and showed broad reactivity to *S. suis* strains from various genetic backgrounds. Although the capsule impaired IgG access to RPS expressed on the bacterial surface, *S. suis* capsule expression is strongly dependent on environmental conditions and varies between *in vitro* and *in vivo* conditions. For example, *S. suis* isolated from mouse nasal-associated lymphoid tissue and CSF at early infection time showed limited CPS expression ^48^, which is in line with observations that CPS biosynthesis genes are downregulated when *S. suis* is cultured in CSF compared to THB ^43^. In addition, 34% of the endocarditis isolates from swine were unencapsulated because of mutations in CPS gene cluster ^49^, but the capsule can recover after *in vivo* passages ^50^. Finally, whilst we only tested the serum samples at a 1:100 dilution because of technical limitations, we observed significantly higher binding of IgG to encapsulated *S. suis* cells in samples from vaccinated animals compared to that of the unvaccinated control group. Therefore, we anticipate that RPS-specific antibodies target the bacteria much stronger during *in vivo* infection. Nevertheless, further studies are needed to clarify the full scope of the role and functionality of the vaccine-induced RPS specific antibodies, e.g. their ability to activate the complement system and induce phagocytosis.

The commensal *S. suis* lineages showed clearly different genetic structures and RPS biosynthesis gene sequences compared to pathogenic lineages. This is of interest, since these findings could imply that vaccination with RPS of pathogenic *S. suis* lineages may likely not affect the commensal non-pathogenic *S. suis* population. If true, this would be beneficial for pig health and reduce the problem of pathogenic strain replacement.

In summary, we identified and characterized the biosynthetic gene cluster and glycosyl linkage patterns of two structural RPS variants from pathogenic *S. suis*. In addition, we demonstrated that the RPS side-chain is important for *S. suis* growth and morphology. The two RPS variants shared a glycan motif that elicited antibodies that were reactive against a broad range of pathogenic *S. suis* strains, which merits future exploration as a component of universal vaccines against the zoonotic pathogen *S. suis*.

## Method

### Identification of *S. suis* RPS biosynthesis gene cluster

The genome sequence of *Streptococcus suis* (*S. suis*) strain P1/7 (NCBI accession: NC_012925.1) was used as the reference. Orthologs of rhamnose-rich polysaccharide (RPS) genes were identified by BLAST (https://blast.ncbi.nlm.nih.gov/Blast.cgi) ^51^. Protein annotation was performed using UniProt (https://www.uniprot.org/) ^52^, InterPro (https://www.ebi.ac.uk/interpro/) ^53^ or Pfam (http://pfam.xfam.org/) ^54^ to identify functional domains. In addition, glycosyltransferases were identified using CAZy (http://www.cazy.org/) ^55^, which is a database for carbohydrate-active enzymes. Genes were visualized by ggplot2 (version 3.3.2) ^56^ and gggenes (version 0.4.0) ^57^ in R (version 4.0.3) ^58,59^. RPS gene cluster alignments were generated with Clinker (version 0.0.23) ^60^ and GC content of RPS cluster was visualized by Artemis (version 18.1.0) ^61^.

### Collection of *S. suis* genomes

A database containing 1,719 publicly-available *S. suis* genomes was constructed previously ^30^. Briefly, 1,800 *S. suis* genomes along with associated metadata were collected from public databases. Genome quality was assessed by QUAST (version 5.0.2) ^62^, any genome with more than 500 contigs, a N50 lower than 10 Kbp, a genome size outside 1.6 – 3.0 Mbp, a GC content outside 40.0 – 42.5% or more than 50 uncalled bases (N’s)/100 Kbp was excluded. Following these quality control measures, 1,719 genomes were retained for analysis in this study.

### Phylogenetic analysis

*S. suis* genomes were annotated by Prokka (version 1.14.0) ^63^ and the results were used as the input for Roary (version 3.12.0) ^64^ to identify the core genome. Core genome sequences were subsequently aligned by MAFFT (version 7.307) ^65^ and sites without single nucleotide polymorphisms (SNPs) were removed by SNP-sites (version 2.5.1) ^66^. The resulting alignment was used to construct a maximum-likelihood phylogenetic tree in IQ-TREE (version 1.6.6) ^67^ with 1,000 bootstrap replicates using the GTR+F+I+G4 model ^68,69^. The final tree was visualized and annotated by iTOL ^70^.

### Conservation analysis of rhamnose and RPS genes

Putative dTDP-rhamnose and RPS biosynthesis genes were screened across the 1,719 genomes by ABRicate (https://github.com/tseemann/abricate) with an initial threshold of 60% coverage and 80% identity. The output was further selected using a more stringent threshold of 90% coverage and 80% identity. Gene conservation was assessed by multiplying coverage and identity and was visualized as heatmaps by iTOL ^70^, together with the phylogenetic tree, sequence type (ST), capsular polysaccharide type (CPS, serotype), host, and geographical location. Only STs and CPS types consisting of more than 10 strains were highlighted.

### Nucleotide variation analysis

The nucleotide sequence of P1/7 whole RPS cluster was screened across the collected genomes by ABRicate (https://github.com/tseemann/abricate, version 1.0.1) using cutoff values of 90% coverage and 80% identity to identify the coordinates of homologous RPS cluster in each genome. These coordinates were used to extract the whole RPS sequences by extract_genes_ABRicate (https://github.com/boasvdp/extract_genes_ABRicate), resulting in 1,350 RPS identified sequences. A more stringent selection was then applied using cutoff values of 95% coverage and 80% identity. Multiple sequence alignment of the selected whole RPS cluster sequences of isolates in pathogenic lineages was made by MAFFT (version 7.471) ^65^ after removing sequence gaps by BioEdit (version 7.0.9.0) ^71^. The aligned sequences were entered into ClustalX (version 2.1) ^72^ to calculate the “column scores”, showing nucleotide site conservation. The variation score was calculated as (100 - column score) and visualized by ggplot2 (version 3.3.2) ^56^ in R (version 4.0.3) ^58,59^.

### Bacterial strains and culture conditions

The principal strains analyzed in this study were *S. suis* strain S10 (ST-1, CPS-2) ^73^, strain 861160 (ST-20, CPS-2) ^74^ and their capsular-deficient mutants S10 ΔCPS ^75^ and 861160 ΔCPS. In addition, *S. suis* strains representing various CPS types and STs were included in the IgG binding assay. A complete list of strains used in this study is provided in Supplementary Table 4. *S. suis* strains were cultured in Todd-Hewitt broth (Oxoid) supplemented with 0.5% yeast extract (Bacto) (THY) at 37°C with gentle shaking (120 rpm). *E. coli* strains for cloning purposes were cultured in Luria-Bertani (LB) media at 37°C with shaking at 200 rpm. When required, antibiotics were added at the following concentrations: chloramphenicol at 10 μg/mL for *E. coli*, kanamycin at 200 μg/mL for *S. suis*, spectinomycin at 100 μg/mL for *S. suis*, and chloramphenicol at 5 μg/mL for *S. suis*.

### Genetic manipulation of *S. suis*

All *S. suis* mutants were constructed by homologous recombination using competence-inducing peptides ^76^. For construction of the 861160 ΔCPS mutant, the fragment for inactivating the *cpsEF* genes was amplified from S10 ΔCPS mutant ^75^ and transformed to 861160 WT strain. To construct *srp* gene deletion mutants, precise in-frame allelic replacement was performed. Briefly, the upstream and downstream regions of targeted *srp* genes were amplified from genomic DNA and ligated to the kanamycin-resistant Janus Cassette ^77^ using overlapping PCR (Phusion™ Hot Start II DNA Polymerase, Thermo Scientific F549S) to construct the knockout fragment, which was subsequently transformed to *S. suis* strains. All mutants were confirmed using PCR and whole genome sequencing.

Genetic complementations of *srp* gene deletion mutants were performed using the plasmid vector pDC123. The *srp* genes were amplified from genomic DNA using primers containing BglII and BamHI restriction sites. Digested PCR products were subsequently ligated to digested plasmid vector using FastDigest Restriction Enzymes (Thermo Scientific FD0083 and FD0054) and T4 DNA Ligase (Roche 10716359001). The ligation products (recombinant plasmid) were verified by PCR and propagated in *E. coli* strains to gain sufficient quantities for complementation. The plasmids were then introduced into *S. suis* mutants using competence-inducing peptides ^76^. The successful complementation of the gene was confirmed by colony PCR and Sanger sequencing. A complete list of primers and plasmids used in this study is provided in Supplementary Table 5.

### Cell wall isolation

Overnight bacterial cultures were diluted 1:20 in fresh THY broth and grown to late exponential phase (O.D._600_ ≈ 0.7). Bacteria were harvested by centrifugation in 4°C and cell walls were isolated using the SDS-boiling procedure, as described before ^22,78^. Isolated cell wall samples were lyophilized and stored at -20°C until further analysis ^32^.

### RPS isolation and purification

*S. suis* RPS was released from the cell wall by mild acid hydrolysis after chemical *N*-acetylation, as described previously ^32^. Isolated RPS was purified by size exclusion chromatography on a BioGel P150 (Bio-Rad 81303) column equilibrated in 0.2 N sodium acetate (NaOAc), pH 3.7, 0.15 M sodium chloride (NaCl). If necessary, the fractions were further purified by ion exchange chromatography on DEAE-Toyopearl column and eluted with NaCl gradients (0 – 0.5 M) ^32^. Total rhamnose and hexose contents were estimated by a modified anthrone assay, as described before ^22,32^.

### Phosphate assay

During chromatography, total phosphate content of each fraction was determined by malachite green method, as described previously ^32^. The total phosphate content of purified RPS was determined by a Malachite green phosphate assay kit (Cayman Chemical 10009325) after hydrolysis and enzymolysis. In detail, purified RPS (equivalent to 700 nmol of rhamnose) was heated in 2 N hydrochloric acid (HCl) at 100°C for 2 h in a total volume of 250 μL. The solution was then cooled on ice and neutralized by sodium hydroxide (NaOH) in the presence of 62.5 mM HEPES pH 7.5. The pH after neutralization was checked by indicator strips. Following this, 50 μL of the acid hydrolyzed sample was incubated overnight at 37°C with 1 U alkaline phosphatase (ThermoFisher 78390), 10 μL 10 × alkaline phosphatase buffer and 1 μL 0.1 M magnesium chloride (MgCl_2_), in a total volume of 100 μL, with continuous rotation. Released phosphate was quantified using the Malachite green phosphate assay kit according to the manufacturer’s instructions.

### Lectin or antibody binding to whole bacteria

Bacterial overnight culture was diluted 1:20 in fresh THY and incubated to mid exponential phase (O.D._600_ = 0.4 – 0.6) at 37°C with gentle shaking. The culture was harvested by centrifugation at 4,000 × g for 10 min. The bacterial pellet was resuspended in phosphate buffered saline supplemented with 0.1% bovine serum albumin (PBS-BSA buffer) and stored at −20°C until further analysis. For analysis, bacterial suspensions were thawed on ice and further diluted to O.D._600_ = 0.048 in PBS-BSA buffer. Subsequently, 12.5 μL bacterial suspension was incubated in a 96-well V bottom plate with the same volume of pig serum or fluorescein-labelled lectin (Vector Laboratories). The mixture was incubated for 20 min at 4°C in the dark, followed by washing using 125 μL PBS-BSA buffer and centrifugation at 4000 × g for 10 min. For antibody binding, secondary staining was performed by resuspending the washed bacterial pellet in 25 μL 10 μg/mL goat anti-pig IgG (Fc):FITC (Bio-Rad AAI41F) and incubating for 20 min at 4°C in the dark, followed by washing once with PBS-BSA buffer. Bacteria were further stained in 50 μL 5 μg/mL Hoechst 33342 (Invitrogen H1399) in PBS for 1 h in the dark at room temperature, followed by washing with PBS-BSA buffer and resuspended in 100 μL 1% formaldehyde (Sigma-Aldrich 252549) in PBS before acquisition on flow cytometer (CytoFLEX, Beckman Coulter Life Sciences).

The final concentrations of lectins or sera after mixing with bacteria were: pig serum at 1:100 dilution, ricinus communis agglutinin I (RCA I, RCA_120_) at 5 μg/mL, soybean agglutinin (SBA) at 5 μg/mL and succinylated wheat germ agglutinin (sWGA) at 0.25 μg/mL.

### RPS biotinylation and coating to beads

RPS biotinylation was conducted by reductive amination using biotin-amine (AxisPharm AP10507). Purified RPS was incubated with biotin-amine and sodium cyanoborohydride (NaCNBH_3_, Sigma-Aldrich 156159) at a molar ratio of 1:600:2500 in 80 mM NaOAc pH 5.5. Dimethyl sulfoxide (DMSO) was employed to increase the solubility of biotin-amine. The reaction was left at room temperature overnight in the dark. The biotinylated RPS was purified by washing with Milli-Q water at least 8 times using centrifugal filter units (Amicon® Ultra Centrifugal Filter, 3 kDa MWCO, Millipore UFC5003) as per the manufacturer’s recommendations (40° fixed angle rotor, 14000 × g, room temperature).

Biotinylated RPS was coated to streptavidin beads (Invitrogen 11205D) by adding 5 × 10^7^ beads (in 20 μL PBS) to 60 μL 0.2 mM RPS and incubating at room temperature for 15 min with shaking. The coated beads were washed three times using PBS and resuspended in 1.25 mL PBS-BSA buffer supplemented with 0.05% Tween_20_ (PBS-BSA-T_20_ buffer). The coating of RPS beads was validated by lectin binding assay and analyzed by flow cytometry (BD FACSCanto II, BD Biosciences).

### Lectin or antibody binding to RPS-coated beads

An amount of 1 × 10^5^ RPS-coated beads or uncoated beads in 12.5 μL PBS-BSA-T_20_ buffer was incubated with the same volume of pig serum or fluorescein labelled lectin (Vector Laboratories) in a 96-well U bottom plate at 4°C for 20 min in the dark with shaking. The beads were washed once using a plate magnet (Invitrogen 12331D). For antibody binding, the secondary staining was performed by resuspending the beads in 25 μL 10 μg/mL goat anti-pig IgG (Fc):FITC (Bio-Rad AAI41F) and incubating for 20 min at 4°C in the dark with shaking, followed by washing once using PBS-BSA-T_20_ buffer. The stained beads were resuspended in 100 μL PBS-BSA-T_20_ buffer and analyzed by flow cytometry (BD FACSCanto II, BD Biosciences or CytoFLEX, Beckman Coulter Life Sciences).

The concentrations of lectins or sera after mixing with beads were the same as those used to stain whole bacteria.

### Flow cytometry data analysis

Data acquired by flow cytometer was analyzed using FlowJo (version 10.10.0). The bacterial population was gated first based on the Hoechst 33342-positive population then using the forward and side scatter plot. The single bacteria population was further selected using the forward-area and forward-height plot. Bacteria stained by only secondary antibody (no pig serum) in bacteria-antibody binding assay or unstrained bacteria in bacteria-lectin binding assay were used as background staining. In contrast, the single beads population was gated based on the forward and side scatter plot. Uncoated beads stained by either lectin or primary serum and secondary antibody were used as background staining.

The antibody level or lectin binding level was represented by delta geometric mean fluorescence intensity (ΔGMFI), which was calculated by subtracting the GMFI of background staining from the GMFI of tested samples.

### Glycosyl composition and linkage analysis

Purified RPS was reduced in 0.1 M ammonium hydroxide (NH_4_OH, Sigma-Aldrich 221228) and 10 mg/mL sodium borohydride (NaBH_4_, Sigma-Aldrich 213462) at room temperature overnight, followed by neutralization using acetic acid (CH₃COOH). Reduced RPS was analyzed by combined gas chromatography-mass spectrometry (GC-MS) in the Complex Carbohydrate Research Center (University of Georgia, Athens, USA) using trimethylsilylated methyl glycoside derivatization for glycosyl composition and using methylation for linkage determination, as described previously ^79^.

### NMR spectroscopy

RPS from *S. suis* were dissolved in D_2_O (0.55 mL). NMR experiments were conducted in 5 mm outer diameter NMR tubes on Bruker NMR spectrometers operating at ^1^H frequencies of 400 or 700 MHz at temperatures of 23°C or 50°C, respectively, using experiments suitable for resonance assignments of glycans ^33,80^. ^1^H NMR chemical shifts were referenced to internal sodium 3-trimethylsilyl-(2,2,3,3-^2^H_4_)-propanoate (*δ*_H_ 0.0), ^13^C chemical shifts were referenced to external dioxane in D_2_O (*δ*_C_ 67.4) and ^31^P chemical shifts were referenced to external 2% H_3_PO_4_ in D_2_O (*δ*_P_ 0.0). Acquired NMR data were processed and analyzed using the TopSpin® software from Bruker.

### Growth curve

Overnight bacterial cultures were diluted in fresh THY without antibiotics to O.D._600_ ≈ 0.4, subsequently further diluted 20-fold and aliquoted to two test tubes, 5 mL/tube, and recorded the O.D._600_ values as time = 0 h. All tubes were incubated at 37°C with gentle shaking. The O.D._600_ value of each tube was measured every 30 min. An extra tube filled with 5 mL fresh THY was used as blank control. Growth curves were made using the average O.D._600_ of the aliquots after subtraction of the blank control.

### Chain length quantification

Stationary phase bacteria from overnight cultures in THY broth were used for Gram staining. Multiple pictures were taken for every strain under 40 × microscope objective. Chain length was quantified by counting the pixels of all bacterial chains in every single picture using ImageJ ^81^, until at least 200 chains were analyzed. The number of pixels was transferred to length (μm) and used for further analysis.

### Scanning electron microscopy (SEM)

Overnight bacterial cultures were diluted 1:10 in fresh THY and incubated to exponential phase (O.D._600_ ≈ 0.4) at 37°C with gentle shaking. Four mL bacterial culture was harvested by centrifugation at 4,000 rpm for 10 min at room temperature. The pellet was resuspended in 1 mL fixative (3% glutaraldehyde/3% paraformaldehyde in 0.1 M phosphate buffer, pH 7.4, Electron Microscopy Sciences 16538-06) and incubated at room temperature for 30 min. Samples were stored at 4°C until analysis.

*S. suis* cultures in fixative were washed with H_2_O, followed by a dehydration with increasing concentrations of alcohol. Samples were incubated with 30%, 50%, 70%, 80%, 90%, 96% and 100% alcohol for 10 min per step at room temperature. After each step, samples were centrifuged at 5,000 rpm for 5 min and supernatant was removed. Samples were adhered to glass slides covered with Poly-L-Lysin and rapidly critically point dried using liquid CO_2_ (Leica Microsystems CPD300, Auto mode, CO_2_ in speed slow, Exchange speed 3 with 14 cycles, Gas out heat slow at speed slow 100%). Subsequently, samples were attached to aluminum stubs (Agar Scientific AGG301) using carbon stickers (Agar Scientific G3347) and sputter coated with 6 nm platinum/palladium coating (Leica Microsystems ACE600). Samples were then imaged using a Zeiss Gemini Sigma 300 FEG SEM at 3 kV using SE2 detector.

### Transmission electron microscopy (TEM)

*S. suis* cultures in fixative were washed with H_2_O for 10 min and incubated for 1 h in osmium tetraoxide (1% in ddH_2_O). Samples were then washed with H_2_O for 10 min and dehydrated with increasing concentrations of alcohol (70%, 70%, 80%, 90%, 90%, 96%, 100% and 100% for 15 min per step). After each step, samples were centrifuged at 5,000 rpm for 5 min and supernatant was removed. Then, samples were impregnated with 1:1 epon (Electron Microscopy Sciences) and propylene oxide for 2 h followed by centrifuging for 10 min at 7,000 rpm and replacing epon/propylene oxide mixture with 100% epon. After incubation overnight, epon was replaced with fresh epon and polymerised at 60°C for 2 days. Samples were sectioned using a diamond knife (Diatome) and ultramicrotome (Leica Microsystems EM UC7) in 60 nm sections and placed on copper grids followed by post staining with uranyl acetate (3.5% in ddH_2_O) and lead citrate (Electron Microscopy Sciences 22410). Samples were then imaged partly using a Talos L120c TEM with 16M BM-CETA camera and partly using a Tecnai T12 TEM with Xarosa camera.

### Lysozyme resistance assay

Knocking out *srp* genes resulted in *S. suis* chain length changes, affecting colony forming units (CFU). We alternatively used an O.D._600_ based strategy to analyze *S. suis* lysozyme resistance. In detail, bacterial overnight cultures were diluted 1:10 in fresh THY and sub-cultured to mid exponential phase (O.D._600_ = 0.4 – 0.6) at 37°C with gentle shaking. All sub-cultures were adjusted to O.D._600_ = 0.4. To harvest the bacteria, 5 mL O.D._600_ = 0.4 culture was centrifuged at 4,000 × g for 5 min at room temperature and washed once using PBS. The bacterial pellet was resuspended in 1 mL PBS. Subsequently, 100 µL bacteria suspension was incubated together with lysozyme from chicken egg white (Sigma-Aldrich L6876) in a 96-well flat bottom plate. The O.D._600_ value was measured immediately to check the input of the bacteria. The plate was incubated at 37°C for 1.5 h with gentle shaking and O.D._600_ value was measured afterwards. Bacteria incubated with PBS (0 µg/mL lysozyme) served as controls. The survival factor was calculated by dividing the O.D._600_ value of the tested samples by the O.D._600_ value of the controls.

### Preparation of glycan-protein conjugates

861160 Δ*srpL* RPS was used to prepare the glycoconjugate. RPS was extracted from the cell wall by mild acid hydrolysis and then purified by size exclusion chromatography, following the method described above. The purified RPS was derivatized at their reducing end with alkyne-hydrazide (Lumiprobe 41770) through reductive amination, using a molar ratio of RPS:hydrazide:NaCNBH_3_ = 1:300:1,000. This reaction was conducted in 100 mM NaOAc at pH 4.5, at room temperature, overnight with gentle shaking. The alkyne-derivatized RPS was subsequently purified by ultracentrifugation, through 25 cycles of washing with Milli-Q water (with each cycle involving approximately a twofold dilution), using centrifugal filter units (Amicon® Ultra Centrifugal Filter, 3 kDa MWCO, Millipore UFC5003) as per the manufacturer’s instructions (40° fixed angle rotor, 14,000 × g, room temperature).

The alkyne-derivatized RPS was then conjugated to CRM_197_-azide (CRM_197_-N_3_) using click chemistry (copper-catalyzed azide-alkyne cycloaddition, CuAAC). Conjugation-ready CRM_197_-N_3_ was ordered from Fina Biosolutions, pre-derivatized with approximately 15 azide groups for reaction with alkyne groups (https://finabio.net/product/crm-azide/). The click reaction was carried out with final concentrations of copper sulfate (Cu_2_SO_4_) at 1 mM, sodium ascorbate at 10 mM, tris(benzyltriazolylmethyl)amine (THPTA) at 1 mM, and aminoguanidine at 10 mM in 30 mM HEPES buffer at pH 8.0. Different ratios of CRM_197_-N_3_ and RPS-alkyne were tested to determine the optimal ratio, which was analyzed via SDS-PAGE. The final chosen molar ratio of azide:alkyne was 1:0.7, corresponding to CRM_197_:RPS = 1:10.5, for glycoconjugate synthesis. The reaction was proceeded at room temperature for 1 h, after which a sample was taken. The sample reaction was stopped by adding Laemmli Sample Buffer supplemented with β-mercaptoethanol (Bio-Rad 1610747) and analyzed by SDS-PAGE (Supplementary Fig. 6a). The conjugate was purified by ultracentrifugation, with 25 cycles of washing in PBS, using centrifugal filter units (Amicon® Ultra Centrifugal Filter, 30 kDa MWCO, Millipore UFC8030) as per the manufacturer’s recommendations (swinging bucket rotor, 4,000 × g, room temperature), followed by sterile filtration through 0.22 μm filters. The filter was washed twice with PBS to maximize recovery. Protein and glycan concentrations were measured using the BCA protein assay kit (Pierce™ BCA Protein Assay Kits, Thermo Scientific 23227) and a modified anthrone assay, respectively.

### Pig immunization and serum collection

CRM_197_-RPS conjugate were adjuvanted with X-Solve, an oil in water emulsion consisting of a 5 to 1 v/v blend of a paraffin-based micro-emulsion and a Vitamin E acetate-based nano-emulsion, mixed in a volume ratio of 1:1 ^39^. Twenty-four three-week-old piglets (Duroc and Yorkshire, both sexes) were divided into two groups: a vaccine group and an unvaccinated control group. All piglets were born and housed on the same farm during the study, with litters evenly distributed across treatment groups. Piglets were immunized intramuscularly twice at a two-week interval (days 0 and 14, corresponding to three-week and five-week of age, respectively) with 2 mL formulation (corresponding to 109 µg carrier protein). Piglets were weaned three days after the first vaccination and then housed in groups. Blood samples were collected on study days 0, 14, and 28 to determine serum antibody levels.

Total serum IgG against RPS and whole bacteria was measured by flow cytometry, and cesarean-derived colostrum-deprived piglet serum (CDCD serum) was used as negative control. In contrast, total serum IgG against the carrier protein was determined by Enzyme-linked immunosorbent assay (ELISA).

### Ethics statements of animal experiment

The Dutch Central Authority for Scientific Procedures on Animals and the Animal Welfare Body of MSD Animal Health approved of this study (Act on Animal Experimentation permit number AVD22100202114853). All researchers performing experimental procedures were qualified to handle the animals in accordance with the Experiments on Animals Act. Housing and management conditions were in line with the EU Directive 2010/63/EU on the protection of animals used for scientific purposes. The health and behavior of the pigs were monitored daily by certified animal caretakers.

### Enzyme-linked immunosorbent assay (ELISA)

Nunc MaxiSorp plates (Thermo Scientific) were coated overnight at 4°C with 0.57 µg/mL CRM_197_ carrier protein in carbonate buffer. After coating, the plates were blocked with 1% BSA in 0.04 M PBS for 1 h at 37°C. After blocking, the plates were washed three times. Serum was added and incubated for 1 h at 37°C. For the detection of total IgG, plates were incubated with 0.1 µg/mL peroxidase labelled goat anti-swine IgG (heavy plus light chain, SeraCare 5220-0363) for 1 h at 37°C. After each incubation step the plates were washed. The plates were developed with TMB as substrate and absorbances were measured at 450 nm.

Every sample was measured in a singular series of eight three-fold dilutions starting with 1:160. A pool of control group sera on day 28 served as background control at 1:160. The antibody titers were defined as the interpolated reciprocal of the dilution corresponding to the O.D. value equal to twice the average background absorbance and are expressed as log_2_ titer.

### Statistical analysis

All data were analyzed and visualized using GraphPad Prism (version 10.2.0). The specific statistical methods are detailed in the figure captions. A *p*-value ≤0.05 was considered statistically significant.

## Contributions

N.M.v.S, C. Schultsz, L.B., A.S., A.A.C.J. and Y.S. conceptualized and designed the project, with valuable input from N.K., J.S.R. and J.D.C.C.. The experiments were conducted by Y.S., G.W., C. Sorieul, T.J.R., J.S.R., M.S., R.G., C.C.D.M., I.M.S., N.N.v.d.W., C.W.K., C.H., P.A. and L.T.. Y.S., N.M.v.S., L.B., C. Schultsz and G.W. wrote the manuscript with input from all authors.

## Acknowledgements

The authors thank Dr. Boas C.L. van der Putten and Dr. Kees C.H. van der Ark (Amsterdam UMC) for bioinformatics analysis, Dr. Edith N. G. Houben, Dr. Yuwei Ding and Dr. Catalin M. Bunduc (Vrije Universiteit Amsterdam) for cell wall and RPS isolation, Dr. Manouk Vrieling (Wageningen Bioveterinary Research) for sharing CDCD serum, Irma Schouten (Amsterdam UMC) for gram staining, Daisy I. Picavet-Havik (Amsterdam UMC) for light microscopy imaging.

