## Supplementary for "A conserved glycan motif overcomes antigenic variation inducing broadly reactive antibodies against the zoonotic pathogen *Streptococcus suis*"

[$] Corresponding author

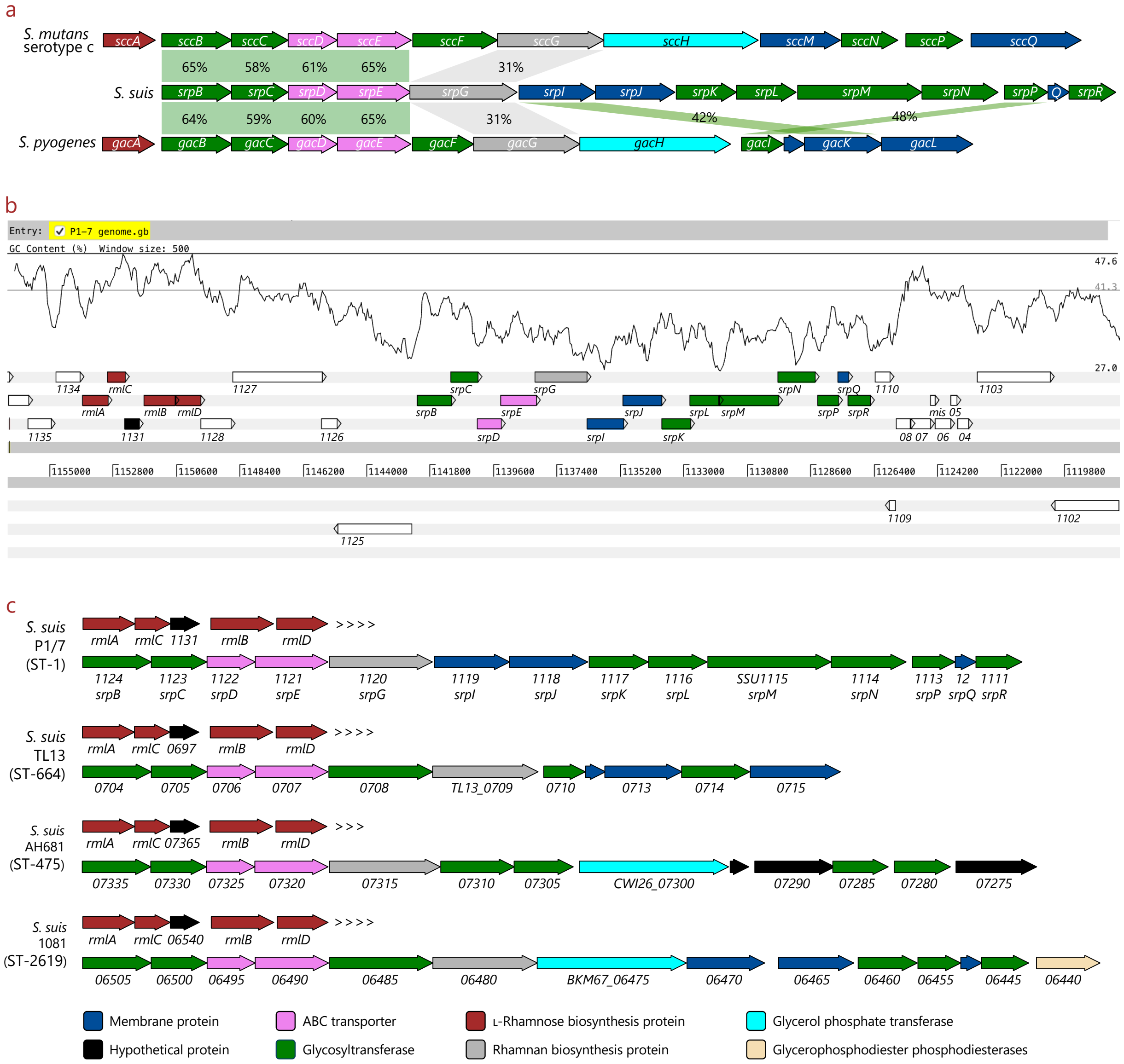


### Supplementary Fig. 1. Homology and genetic structures of the *S. suis* RPS biosynthetic gene cluster compared to other streptococcal species and non-pathogenic *S. suis* lineages

**a**, Comparison of RPS biosynthesis gene clusters of *S. suis* P1/7 with *S. mutans* serotype c strain UA159, and *S. pyogenes* strain MGAS5005. RPS of *S. mutans* serotype c and *S. pyogenes* were also known as *S. mutans* serotype c carbohydrate (SCC) and group A carbohydrate (GAC), respectively. The RPS genes of *S. suis* were named in accordance with their homologs in *S. mutans* and *S. pyogenes*. The *S. mutans* SCC gene cluster represents the gene locus *smu824* – *smu835* in strain UA159 (NCBI accession: NC_004350.2); The *S. suis* RPS gene cluster represents the gene locus *SSU1124* – *SSU1111* in strain P1/7 (NCBI accession: NC_012925.1); The *S. pyogenes* GAC gene cluster represents the gene locus *spy0602* – *spy0613* in strain MGAS5005 (NCBI accession: NC_007297.2). The arrows are scaled according to gene size, with each function represented by a different color. The alignment was created using amino acid sequences, highlighting regions with over 30% identity. **b**, GC content of *S. suis* RPS gene cluster as visualized using Artemis. **c**, Examples of *S. suis* RPS gene clusters with different genetic structures compared to strain P1/7. The three visualized RPS biosynthesis gene clusters were identified in strains from the non-pathogenic lineages. The arrows are scaled according to gene size, with each predicted function represented by a different color. NCBI accessions of the genome: P1/7, NC_012925.1; TL13, NC_021213.1; AH681, NZ_CP025043.1; 1081, NZ_CP017667.1; ST, sequence type.

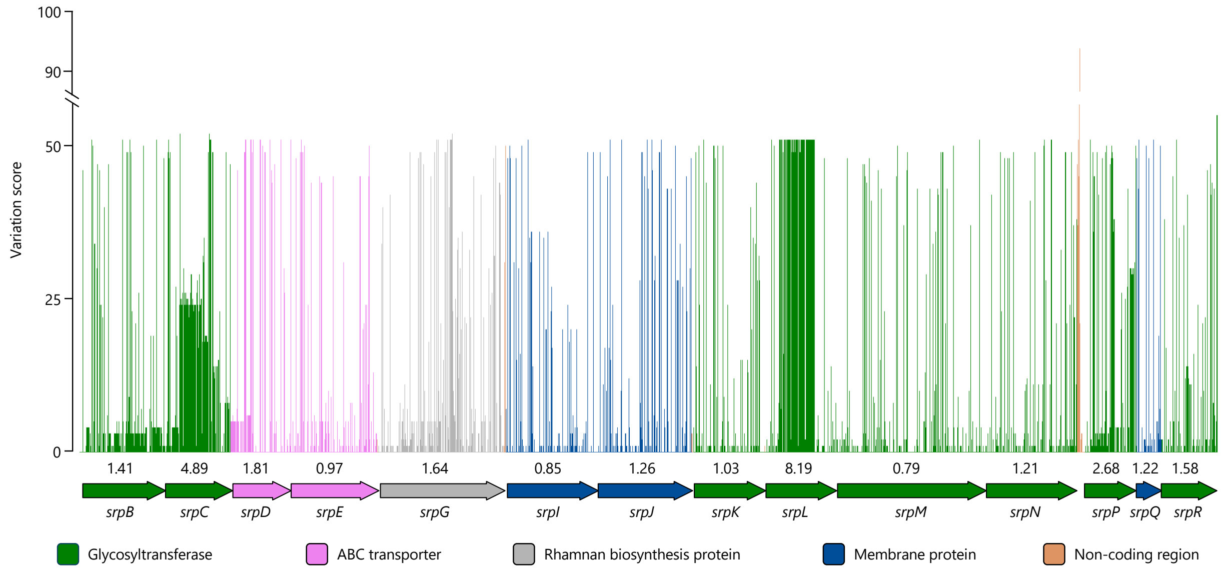


Supplementary Fig. 2. Nucleotide variation within the RPS cluster in strains from the pathogenic *S. suis* lineages

Schematic representation of the nucleotide variation at each position within the RPS cluster of pathogenic *S. suis* isolates in comparison to strain P1/7. The variation score of each nucleotide site was calculated using ClustalX. The mean value of the variation score of each gene is shown below the bar chart. The glycosyltransferase encoding genes *srpC* and *srpL* are more variable than other RPS genes.

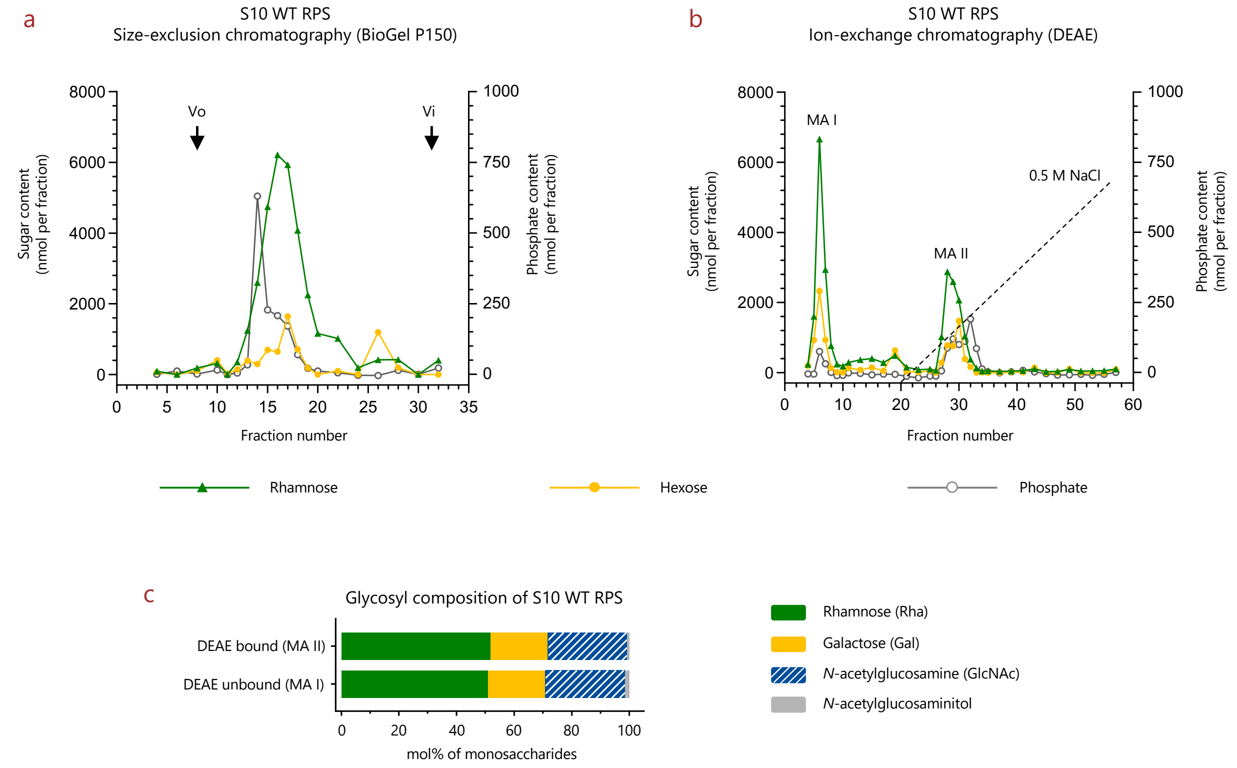


Supplementary Fig. 3. Glycosyl composition analysis of S10 wild-type RPS released from cell wall by chemical treatments and purified by size-exclusion and ion-exchange chromatography

**a**, Size-exclusion chromatography profile of wild-type RPS released from *S. suis* S10 cell wall by mild acid hydrolysis. Fractions chromatographed by BioGel P150 were analyzed for rhamnose and hexose contents by a modified anthrone assay, phosphate by malachite green assay. WT, wild-type. **b**, Ion-exchange chromatography of the S10 WT RPS released by mild acid hydrolysis. Fractions containing the RPS material (from **a**) were pooled, concentrated, desalted by centrifugal filter units, loaded onto DEAE-Toyopearl and eluted with a NaCl gradient (0 – 0.5 M). Fractions were analyzed for rhamnose and hexose content by anthrone assay and total phosphate content by malachite green assay. DEAE unbound fractions (MA I) and DEAE bound fraction (MA II) were submitted to GC-MS for further glycosyl composition and linkage analysis. Vo, void volume; Vi, included volume. **c**, Glycosyl composition analysis by GC-MS of TMS derivatives of methyl glycosides of the two fractions of S10 WT RPS. *N*-acetylglucosaminitol is the reducing end of mild acid hydrolysis. The original data was shown on Supplementary Table 2.

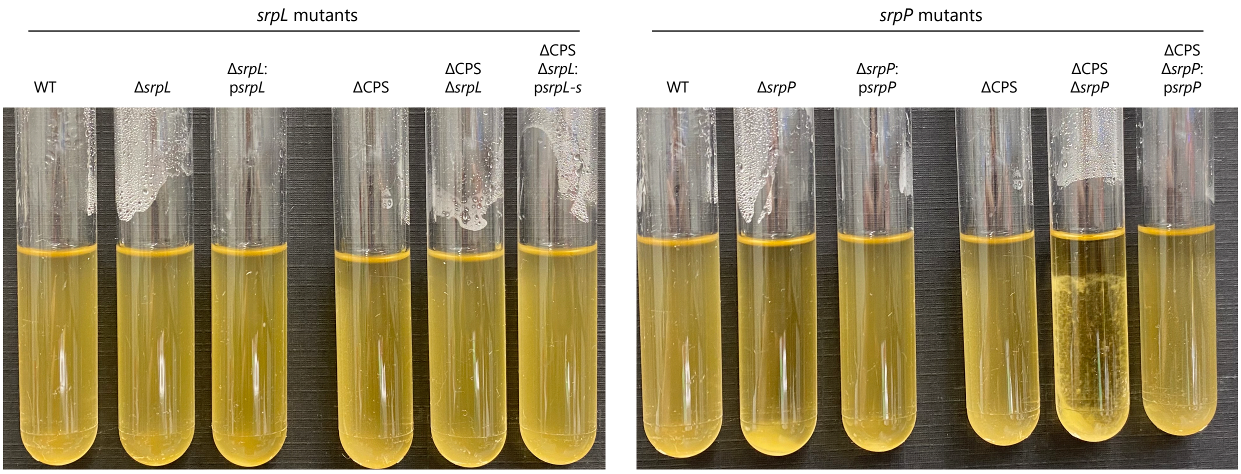


Supplementary Fig. 4. Self-aggregation phenotype of *S. suis* RPS gene deletion mutants

Aggregation phenotype of *S. suis* S10 wild-type and Δ*srpL* (left) and Δ*srpP* (right) mutants and complemented strains. Bacteria were grown in THY broth overnight with gentle shaking. The experiment was performed independently three times and yielded similar results. WT, wild-type.

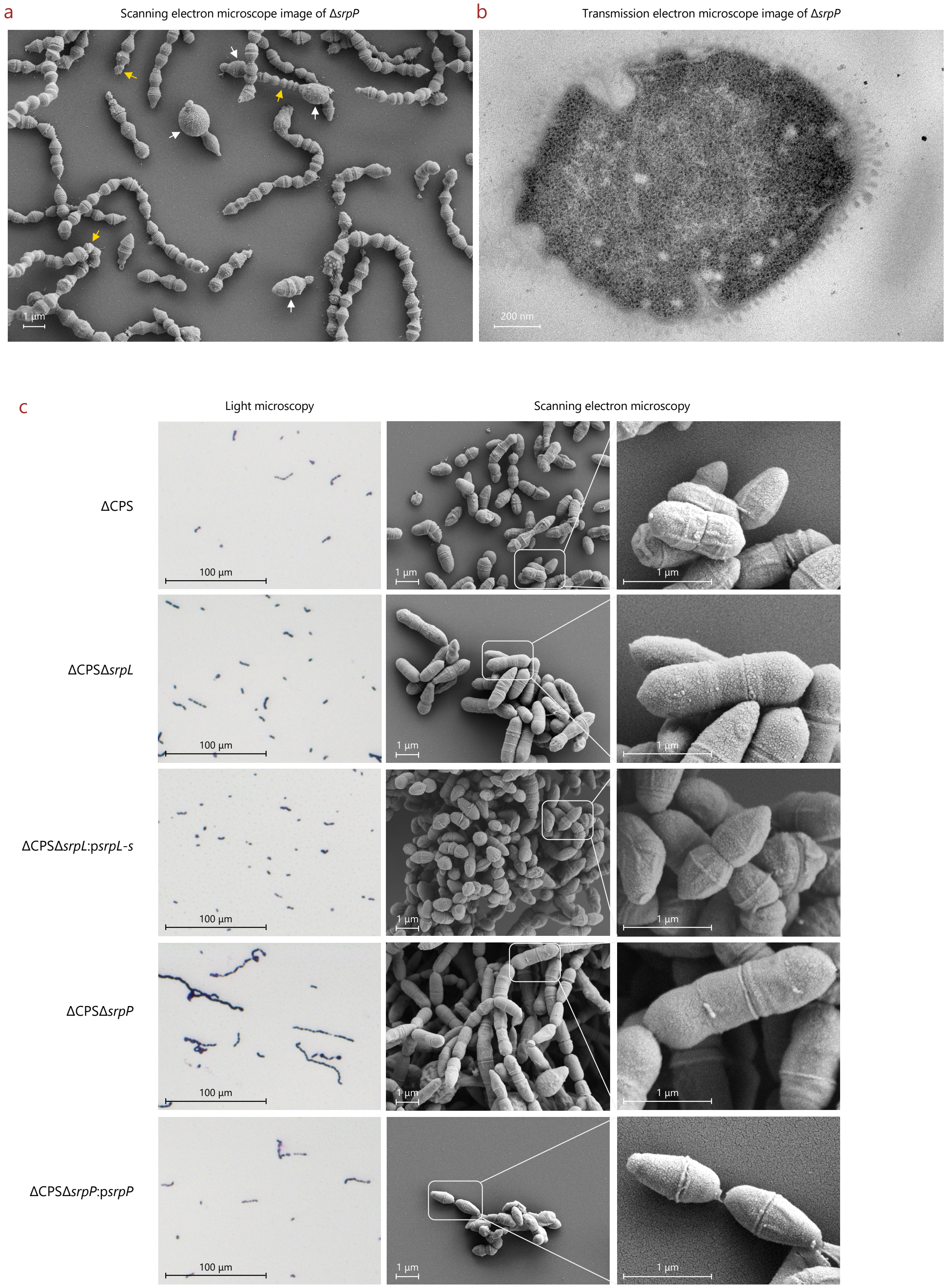


### Supplementary Fig. 5. Representative microscopy images of RPS gene deletion mutants

**a** and **b**, Representative electron microscope images of encapsulated S10 Δ*srpP*. **a**, Scanning electron microscope images with white arrows denote swollen and large bacterial cells; the yellow arrows denote small blebs or fractional bacterial cells. **b**, Transmission electron microscope images of a representative swollen bacterial cell. **c**, Representative microscopy images of capsule deficient S10 and RPS gene deletion mutants. Light microscopy images were taken on

Gram staining slides of stationary phase bacteria from overnight culture in THY, whereas electron microscopy images were taken on exponential phase bacteria ($O.D._{600} \approx 0.4$ in THY).

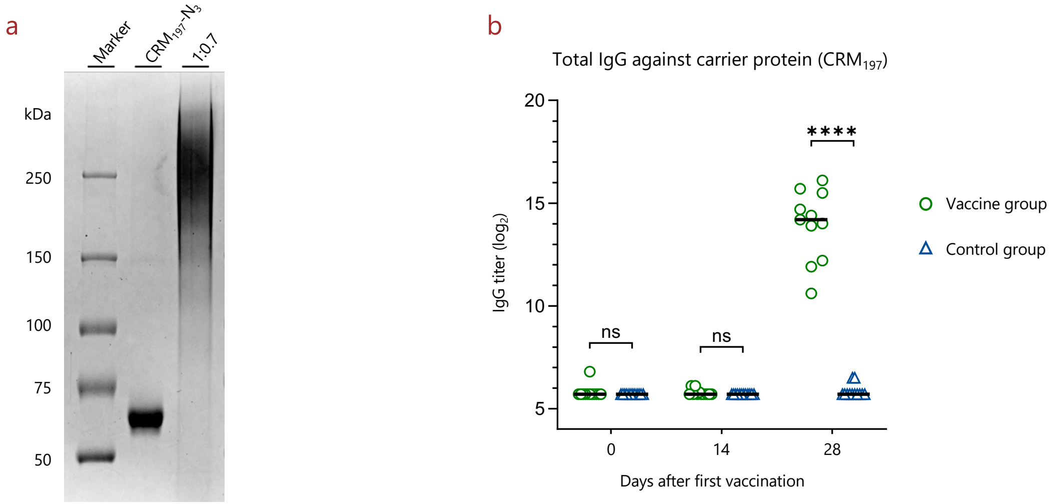


Supplementary Fig. 6. Analysis of the RPS glycoconjugate and immune response against the carrier protein

**a**, SDS-PAGE analysis of the RPS-$CRM_{197}$ conjugate. Three µg of each sample was loaded onto a gel (7.5% acrylamide) for SDS-PAGE analysis. A molar ratio of 1 azide ($N_3$) to 0.7 RPS-alkyne corresponding to 1 $CRM_{197}$-$N_3$ to 10.5 RPS-alkyne was used to make the conjugate. No remaining free protein could be observed. **b**, RPS-$CRM_{197}$ conjugate induced antibodies against the carrier protein on day 28. Total IgG titer was determined by ELISA. The line represents the median value. *P* values were calculated by Mann-Whitney test with Holm-Šídák's multiple comparisons test. ****$p \leq 0.0001$; ns, not significant.

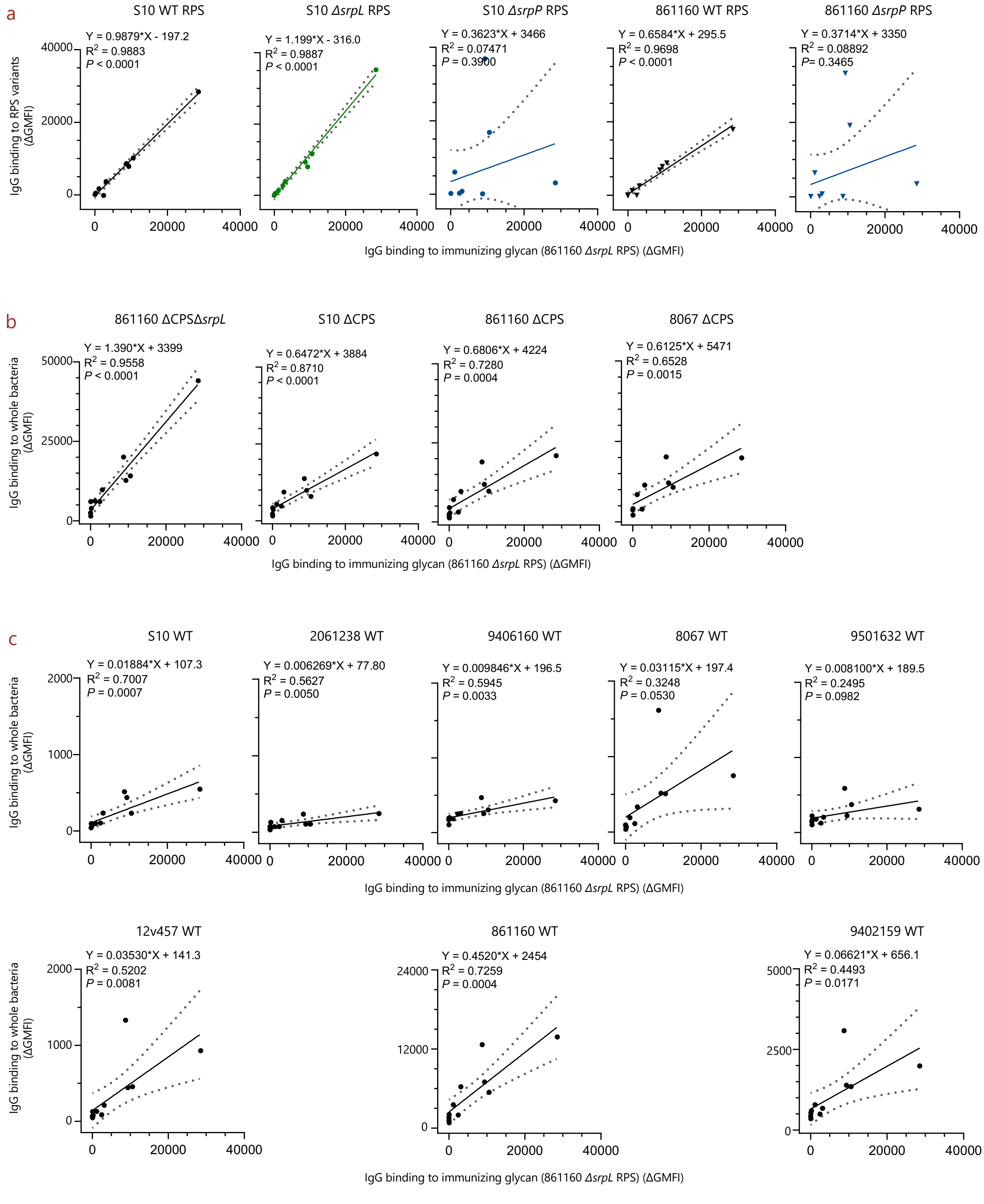


Supplementary Fig. 7. Correlation plots for IgG binding to immunizing glycan and different antigens or bacterial strains

**a**, Correlations between IgG binding to the immunizing glycan and different RPS variants; **b**, Correlations between IgG binding to the immunizing glycan and different *S. suis* capsule-deficient strains; **c**, Correlations of IgG binding to the immunizing glycan and encapsulated wild-type strains from different genetic backgrounds. Correlations were shown using simple linear regression with 95% confidence interval. WT, wild-type.

Supplementary Table 1. *S. suis* putative RPS biosynthetic genes and their predicted functions

| ***S. suis* strain[1]** | | | | **Gene name** | **Predicted function of the encoded protein[2]** |
|---|---|---|---|---|---|
| **P1/7** | **05ZYH33** | **ZY05719** | **861160** | | |
| Rhamnose biosynthesis genes | | | | | |
| *SSU1133* | *SSU05_1301* | *06205* | *RS06805* | *rmlA* | Glucose-1-phosphate thymidylyltransferase. |
| *SSU1132* | *SSU05_1300* | *06200* | *RS06800* | *rmlC* | dTDP-4-dehydrorhamnose 3,5-epimerase. |
| *SSU1131* | *SSU05_1299* | *06195* | *RS06795* | | Hypothetical protein. |
| *SSU1130* | *SSU05_1298* | *06190* | *RS06790* | *rmlB* | dTDP-glucose 4,6-dehydratase. |
| *SSU1129* | *SSU05_1296* | *06185* | *RS06785* | *rmlD* | dTDP-4-dehydrorhamnose reductase. |
| RPS biosynthesis genes | | | | | |
| *SSU1124* | *SSU05_1289* | *06160* | *RS06760* | *srpB* | Glycosyltransferase. 47/381 (65%) amino acid identity to *S. pyogenes* GacB. |
| *SSU1123* | *SSU05_1288* | *06155* | *RS06755* | *srpC* | Glycosyltransferase. 183/308 (59%) amino acid identity to *S. pyogenes* GacC. |
| *SSU1122* | *SSU05_1287* | *06150* | *RS06750* | *srpD* | ABC transporter. 160/268 (60%) amino acid identity to *S. pyogenes* GacD. |
| *SSU1121* | *SSU05_1286* | *06145* | *RS06745* | *srpE* | ABC transporter. 261/398 (66%) amino acid identity to *S. pyogenes* GacE. |
| *SSU1120* | *SSU05_1285* | *06140* | *RS06740* | *srpG* | Rhamnan backbone synthesis protein. 185/584 (32%) amino acid identity to *S. pyogenes* GacG. |
| *SSU1119* | *SSU05_1284* | *06135* | *RS06735* | *srpI* | Membrane protein. 180/403 (45%) amino acid identity to *S. pyogenes* GacK. |
| *SSU1118* | *SSU05_1282* | *06130* | *RS06730* | *srpJ* | Membrane protein with a DUF2142 domain. |
| *SSU1117* | *SSU05_1281* | *06125* | *RS06725* | *srpK* | Glycosyltransferase with a GTF[3] 8 domain |
| *SSU1116* | *SSU05_1280* | *06120* | *RS06720* | *srpL* | Glycosyltransferase with a GTF 2 domain. |
| *SSU1115* | *SSU05_1279* | *06115* | *RS06715* | *srpM* | Glycosyltransferase with rhamnan synthesis F domain and a GTF 1 domain. |
| *SSU1114* | *SSU05_1278* | *06110* | *RS06710* | *srpN* | Glycosyltransferase with a GTF 4 domain and a GTF 1 domain. |
| *SSU1113* | *SSU05_1277* | *06105* | *RS06705* | *srpP* | Glycosyltransferase. 113/233 (48%) amino acid identity to *S. pyogenes* GacI. |
| *SSU1112* | *SSU05_1276* | *06100* | *RS06700* | *srpQ* | Membrane protein with a DUF2304 domain. |
| *SSU1111* | *SSU05_1275* | *06095* | *RS06695* | *srpR* | Glycosyltransferase. 121/258 (47%) amino acid identity to *L. lactis* WpsC. |
| *gacO*, *gbcO* and *rgpG* homolog | | | | | |
| *SSU1672* | *SSU05_1877* | *08895* | *RS08935* | *srpO* | UDP-*N*-acetylglucosamine-1-phosphate: lipid phosphate transferase. |

[1] NCBI accessions of *S. suis* genome: P1/7, NC_012925.1; 05ZYH33, CP000407.1; ZY05719, NZ_CP007497.1; 861160, NZ_LR738722.1.

[2] Protein similarities between P1/7 and *S. pyogenes* strain MGAS5005 (NCBI accession: NC_007297.2), or *Lactococcus lactis* subsp. cremoris strain MG1363 (NCBI accession: AM406671.1).

[3] GTF, Glycosyltransferases family.

Supplementary Table 2. Glycosyl linkage analysis by GC-MS of partially methylated alditol acetate derivatives

| Glycosyl Linkage Residue | Relative Area Percentage (%)[1] | | | | | | |
|---|---|---|---|---|---|---|---|
| | S10 | | | | 861160[2] | | |
| | WT | | | | | | |
| | MA I[3] | MA II | Δ*srpL* | Δ*srpP* | WT | Δ*srpL* | Δ*srpP* |
| Terminal Rhamnopyranosyl Residue (t-Rha*p*) | 0 | 0 | 1.7 | 0.5 | 1.0 | 2.1 | 2.4 |
| 2-linked Rhamnopyranosyl Residue (2-Rha*p*) | 18.8 | 12.4 | 34.9 | 66.7 | 25.5 | 39.5 | 61.6 |
| 3-linked Rhamnopyranosyl Residue (3-Rha*p*) | 5.3 | 3.3 | 6.2 | 13.6 | 7.1 | 6.6 | 13.6 |
| 2,3-linked Rhamnopyranosyl Residue (2,3-Rha*p*) | 10.2 | 11.2 | 14.1 | 0 | 9.0 | 11.4 | 0.8 |
| 2,4-linked Rhamnopyranosyl Residue (2,4-Rhap) | 12.8 | 14 | 0.3 | 0.2 | 14 | 0.2 | 0.4 |
| 3,4-Linked Rhamnopyranosyl Residue (3,4-Rha*p*) | | | 0 | 0 | 0 | 0 | 0.5 |
| 2,3,4-linked Rhamnopyranosyl Residue (2,3,4-Rha*p*) | 0.1 | 0.2 | 0.1 | 0 | 0.2 | 0.1 | 0.1 |
| Terminal Glucopyranosyl Residue (t-Glc*p*) | | | 0.1 | 0 | 14.3 | 0.2 | 0.5 |
| 2-linked Glucopyranosyl Residue (2-Glc*p*) | 0.1 | 0.2 | | | | | |
| 6-linked Glucopyranosyl Residue (6-Glc*p*) | 0.1 | 0.2 | | | 0.2 | 0.1 | 0.2 |
| 2,6-linked Glucopyranosyl Residue (2,6-Glc*p*) | 0.1 | 0.3 | | | | | |
| 3-linked Mannopyranosyl Residue (3-Man*p*) | 0.3 | 0.5 | | | | | |
| 6-linked Mannopyranosyl Residue (6-Man*p*) | 0.1 | 0.2 | | | | | |
| Terminal Galactopyranosyl Residue (t-Gal*p*) | 23.9 | 22.6 | 19.1 | 0 | 15.8 | 19.5 | 0 |
| 3-Linked Galactopyranosyl Residue (3-Gal*p*) | | | | | 0.2 | 0 | 0 |
| 4-Linked Galactopyranosyl Residue (4-Gal*p*) | | | 0.1 | 0 | 0.2 | 0.3 | 0.2 |
| 6-linked Galactopyranosyl Residue (6-Gal*p*) | 0.1 | 0.3 | | | 0.1 | 0.1 | 0 |
| 3,4-linked Galactopyranosyl Residue (3,4-Gal*p*) | 0 | 0 | | | | | |
| Terminal 2-Acetamido-2-deoxy-glucopyranosyl Residue (t-GlcNAc) | 2.0 | 2.7 | 0.6 | 0.9 | 1.1 | 1.5 | 2.0 |
| 3-Linked 2-Acetamido-2-deoxy-glucopyranosyl Residue (3-GlcNAc) | 13.1 | 16.6 | 11.8 | 0 | 4.7 | 7.8 | 0 |
| 6-Linked 2-Acetamido-2-deoxy-glucopyranosyl Residue (6-GlcNAc) | 12.0 | 14.7 | 9.7 | 16.6 | 4.0 | 6.7 | 11.0 |
| 4-Linked 2-Acetamido-2-deoxy-glucopyranositol Residue (4-GlcNAcitol)[4] | 0.9 | 0.7 | 0.9 | 1.6 | 1.3 | 1.6 | 3.6 |
| 4-linked 2,5-anhydromannitol | 0 | 0 | | | | | |
| Total | 99.9 | 100.1 | 100 | 100.1 | 100 | 99.9 | 99.9 |

[1] All RPS samples were released from cell wall by mild acid hydrolysis after chemical *N*-acetylation.

[2] 4-Linked Glucopyranosyl Residue (4-Glc*p*) was also detected in 861160 RPS samples, but that was attributed to unexpected contaminations.

[3] MA I is DEAE unbound sample, MA II is DEAE bound RPS sample.

[4] 4-GlcNAcitol is the reducing end of RPS released by mild acid hydrolysis after chemical *N*-acetylation.

Supplementary Table 3. Phosphate content of purified RPS

| | Phosphate content (nmol) normalized to 50 nmol rhamnose | |
|---|---|---|
| | S10 | 861160 |
| WT | 3.19 | 3.35 |
| Δ*srpL* | -0.20 | -0.05 |
| Δ*srpP* | 0.00 | -0.06 |

Phosphate concentration was measured by malachite green assay following hydrolyzation with hydrochloric acid and digestion with alkaline phosphatase. Rhamnose content was determined by a modified anthrone assay, as described in methods. Data are expressed as nmol of phosphate normalized to 50 nmol rhamnose in the sample.

Supplementary Table 4. Bacterial strains and plasmids used in this study

| **Strain or plasmid** | **Description**[a] | **Reference** |
|---|---|---|
| *Streptococcus suis* wild-type strain | | |
| 9402159 | *S. suis* strain isolated from a meningitis pig in the Netherlands in 2006. Capsular type 1, sequence type 13, clonal complex 13. | 1 |
| P1/7 | *S. suis* reference strain isolated from a pig dying with meningitis in the United Kingdom. Capsular type 2, sequence type 1, clonal complex 1. | 2 |
| S10 | Virulent *S. suis* strain which was originally isolated from the tonsils of a healthy pig at a slaughter in the Netherlands in 1992. Also known as strain 10. Capsular type 2, sequence type 1, clonal complex 1. | 3 |
| 2061238 | *S. suis* strain isolated from a human patient with meningitis in the Netherlands in 2006. Capsular type 2, sequence type 146, clonal complex 1. | 1 |
| 861160 | *S. suis* strain isolated from a human patient with meningitis in the Netherlands in 1986. Capsular type 2, sequence type 20, clonal complex 20. | 4 |
| 12v457 | *S. suis* strain isolated from a meningitis pig in the Netherlands in 2006. Capsular type 7, sequence type 29, clonal complex 29. | 1 |
| 9501632 | *S. suis* strain isolated from a meningitis pig in the Netherlands in 2006. Capsular type 8, sequence type 87, clonal complex 16. | 1 |
| 9406160 | *S. suis* strain isolated from a meningitis pig in the Netherlands in 2006. Capsular type 9, sequence type 16, clonal complex 16. | 1 |
| 8067 | *S. suis* strain isolated from a meningitis pig in the Netherlands in 1996. Capsular type 9, sequence type 136, clonal complex 16. | 4 |
| *Streptococcus suis* gene deletion and complementation mutant | | |
| S10 Δ*srpL* | *srpL* deletion mutant in the S10 background by gene replacement with the kanamycin resistance Janus cassette[5]. $Kan^R$ | This study |
| S10 Δ*srpL*:p*srpL* | S10 *ΔsrpL* complemented with plasmid p*srpL* carrying S10 *srpL*. $Kan^R$, $Chl^R$ | This study |
| S10 Δ*srpP* | *srpP* deletion mutant in the S10 background by gene replacement with the kanamycin resistance Janus cassette[5]. $Kan^R$ | This study |
| S10 Δ*srpP*:p*srpP* | S10 *ΔsrpP* complemented with plasmid p*srpP* carrying S10 *srpP*. $Kan^R$, $Chl^R$ | This study |
| S10 ΔCPS | Capsule negative mutant of S10. Capsule biosynthesis genes *cpsEF* were inactivated by the insertion of a spectinomycin resistance cassette. $Spc^R$ | 6 |
| S10 ΔCPSΔ*srpL* | *srpL* deletion mutant in the S10 ΔCPS background by gene replacement with the kanamycin resistance Janus cassette[5]. $Spc^R$, $Kan^R$ | This study |
| S10 ΔCPSΔ*srpL*:p*srpL-s* | S10 ΔCPSΔ*srpL* homologous (self) complementary strain (complemented with plasmid p*srpL* carrying S10 *srpL*). $Spc^R$, $Kan^R$, $Chl^R$ | This study |
| S10 ΔCPSΔ*srpL*:p*srpL-c* | S10 ΔCPSΔ*srpL* heterologous (cross) complementary strain (complemented with plasmid p*srpL* carrying 861160 *srpL*). $Spc^R$, $Kan^R$, $Chl^R$ | This study |
| S10 ΔCPSΔ*srpP* | *srpP* deletion mutant in the S10 ΔCPS background by gene replacement with the kanamycin resistance Janus cassette[5]. $Spc^R$, $Kan^R$ | This study |
| S10 ΔCPS*ΔsrpP*:p*srpP* | S10 ΔCPSΔ*srpP* complemented with plasmid p*srpP* carrying S10 *srpP*. $Spc^R$, $Kan^R$, $Chl^R$ | This study |
| 861160 ΔCPS | Capsule negative mutant of 861160. Capsule biosynthesis genes *cpsEF* were inactivated by the insertion of a spectinomycin resistance cassette. $Spc^R$ | This study |
| 861160 ΔCPSΔ*srpL* | *srpL* deletion mutant in the 861160 ΔCPS background by gene replacement with the kanamycin resistance Janus cassette[5]. $Spc^R$, $Kan^R$ | This study |

| | | |
|---|---|---|
| 861160 ΔCPSΔ*srpL*:p*srpL-s* | 861160 ΔCPSΔ*srpL* homologous (self) complementary strain (complemented with plasmid p*srpL* carrying 861160 *srpL*). Spc$^R$, Kan$^R$, Chl$^R$ | This study |
| 861160 ΔCPSΔ*srpL*:p*srpL-c* | 861160 ΔCPSΔ*srpL* heterologous (cross) complementary strain (complemented with plasmid p*srpL* carrying S10 *srpL*). Spc$^R$, Kan$^R$, Chl$^R$ | This study |
| 861160 ΔCPSΔ*srpP* | *srpP* deletion mutant in the 861160 ΔCPS background by gene replacement with the kanamycin resistance Janus cassette[5]. Spc$^R$, Kan$^R$ | This study |
| 861160 ΔCPSΔ*srpP*:p*srpP* | 861160 ΔCPSΔ*srpP* complemented with plasmid p*srpP* carrying 861160 *srpP*. Spc$^R$, Kan$^R$, Chl$^R$ | This study |
| 8067 ΔCPS | Capsule negative mutant of 8067. Capsule biosynthesis gene *cpsE* was inactivated insertion of spectinomycin resistance cassette. Spc$^R$ | [7] |
| *Escherichia coli* strain | | |
| MC1061 | *E. coli* cells used for cloning. | |
| Top10 | *E. coli* cells used for cloning. | |
| Plasmid | | |
| pDC123 | Complementation vector with a chloramphenicol resistance cassette. Chl$^R$ | [8] |
| p*srpL* | pDC123 derived plasmid expressing *srpL* either from S10 or 861160. Chl$^R$ | This study |
| p*srpP* | pDC123 derived plasmid expressing *srpP* either from S10 or 861160. Chl$^R$ | This study |

[a] Antibiotic resistance markers: Kan$^R$, kanamycin; Spc$^R$, spectinomycin; Chl$^R$, chloramphenicol

Supplementary Table 5. Primers for genetic manipulation

| **Primer** | **Sequence**[a] | **Genetic manipulation** |
|---|---|---|
| CD_1116F/Rcomp | cttggaatgtgcccagaaccgac | S10 and 861160 *srpL* deletion with a kanamycin resistance Janus cassette |
| CD_1116R/Fcomp | cacccatggaaattccccaaaagtc | |
| CD_1116KanR/Fcomp | cattatccattaaaaatcaaacGGacctagactcctttttgtaatctaattttc | |
| CD_1116KanF/Rcomp | ctaaacgtccaaaagcataaGGtaatagaaagtgattggtaacactg | |
| CD_1116KanR/Fcomp1 | cagtgttaccaatcactttctattaCCttatgcttttggacgtttag | |
| CD_1116KanF/Rcomp2 | gaaaattagattacaaaaaggagtctaggtCCgtttgatttttaatggataatg | |
| CD_1116F/Rcomp_WT | gttattatcccagcgtacaatg | Verification of *srpL* |
| CD_1116R/Fcomp_WT | caaaagtttgttttgaataaatgttg | |
| YS_1116-BglII_Fwd | GCGTAAGATCttatctacttttataaattttcaaaagtttgttttgaataaatgttg | Construction of *psrpL* |
| YS_1116-BamHI_Rvs | CGTCTGGATCCgtgaagttttccgttattatcccagc | |
| CD_1113F/Rcomp | caaggagcaattgctgcctcatgtc | S10 and 861160 *srpP* deletion with a kanamycin resistance Janus cassette |
| CD_1113R/Fcomp | gtcgtctcagattcctcatggatacg | |
| CD_1113KanR/Fcomp | cattatccattaaaaatcaaacGGtttttctctctaaaactagaattc | |
| CD_1113KanF/Rcomp | ctaaacgtccaaaagcataaGGAAatgactatacaagcattagctatg | |
| CD_1113KanR/Fcomp1 | catagctaatgcttgtatagtcatTTCCttatgcttttggacgtttag | |
| CD_1113KanF/Rcomp2 | gaattctagttttagagagaaaaaCCgtttgatttttaatggataatg | |
| CD_1113F/Rcomp_WT | gtattaatgattatccctgcctac | Verification of *srpP* |
| CD_1113R/Fcomp_WT | tccttcataagcgaagcaatc | |
| YS_1113-BglII_Fwd | GCGTAAGATCttattccccctccttcataagcgaag | Construction of *psrpP* |
| YS_1113-BamHI_Rvs | CGTCTGGATCCatgaaagtattaatgattatccctgcctacaatgaag | |
| YS_pDC123_Fwd | cgaaacgctaaagcctttcgg | Primers for pDC123 |
| YS_pDC123_Rvs | ggaattgtcagataggcctaatgac | |
| YS_cps2E/F_Fwd | ctcgtatgccggttgaatttgag | Amplification of *cps2EF* mutation construct from S10 ΔCPS |
| YS_cps2E/F_Rvs | ccagtatttcgctccttcttcc | |

[a] Restriction sites are underlined.
